# Diversity-driven biochemical survey reveals dimeric structural origin of rubisco

**DOI:** 10.1101/2025.11.05.686826

**Authors:** Alexander J. Kehl, Leah Taylor-Kearney, Alexander L. Jaffe, Jose Henrique Pereira, Jennifer Lee, Michal Hammel, Lucas M. Waldburger, Caroline Yeow, Luis Valentin-Alvarado, Paul D. Adams, Jillian F. Banfield, Justin B. Siegel, Noam Prywes, Patrick M. Shih

**Affiliations:** Biophysics Graduate Group, University of California, Davis, Davis, CA, USA; Environmental Genomics and Systems Biology Division, Lawrence Berkeley National Laboratory, Berkeley, CA 94720, USA; Feedstocks Division, Joint BioEnergy Institute, Emeryville, CA, USA; Department of Plant and Microbial Biology, University of California, Berkeley, Berkeley, CA 94720, USA; Department of Earth System Science, Stanford University, Stanford, CA USA; Molecular Biophysics and Integrated Bioimaging Division, Lawrence Berkeley National Laboratory, Berkeley, CA, USA; Genome Center, University of California Davis, Davis, CA, USA; Department of Bioengineering, University of California, Berkeley, California, USA; Biological Systems & Engineering Division, Lawrence Berkeley National Laboratory, Berkeley, CA 94720, USA; Department of Electrical Engineering and Computer Science, University of California, Berkeley, California, USA; Department of Biochemistry and Molecular Biology, Biomedicine Discovery Institute, Monash University, Clayton, Australia; Department of Bioengineering, University of California, Berkeley, CA, 94720, USA; Innovative Genomics Institute, University of California, Berkeley, Berkeley, CA, USA; Department of Earth and Planetary Science, University of California, Berkeley, Berkeley, CA, USA; Department of Environmental Science, Policy, and Management, University of California, Berkeley, Berkeley, CA, USA; Department of Chemistry, University of California-Davis, Davis, California 95616, USA; Department of Biochemistry and Molecular Medicine, University of California-Davis, Davis, California 95616, USA; Department of Biochemistry, University of Cambridge, Cambridge, UK

## Abstract

Rubisco is the entry point of nearly all organic carbon into the biosphere and is present in all domains of life. Despite its global importance, biochemical studies of this enzyme superfamily have been limited to a relatively narrow set of subclades. Recent advances in metagenomics have dramatically reshaped our understanding of both microbial and rubisco diversity; however, biochemical characterization of these sequences has not kept pace with the exponential growth in sequence data. To better survey the functional and structural diversity of rubisco, we systematically sampled and synthesized a library of diverse rubisco sequences with an emphasis on clades that have previously not been characterized. Our updated phylogenetic analysis reveals that many deep-branching rubiscos assemble as dimers, supporting a dimeric origin for the superfamily — in contrast to the ecologically dominant hexadecameric form I. Additionally, we discover and structurally characterize the largest rubisco described to date, originating from a cryptic, early-branching subclade with novel structural folds that have previously not been observed in the rubisco superfamily. By integrating biochemical data with an updated phylogenetic framework, we propose a revised nomenclature for the rubisco protein family that reflects current insights and will better accommodate future discoveries.

## Introduction

Ribulose-1,5-bisphosphate carboxylase/oxygenase (rubisco) is the primary carboxylating enzyme on earth^1,2^, facilitating the conversion of atmospheric carbon dioxide (CO_2_) into the building blocks of life in autotrophic organisms like plants, algae, and bacteria^3^. Rubisco evolved at least three billion years ago, and today is found in eukaryotes, bacteria, and archaea^4^. All rubisco sequences form a single clade that is distinct from rubisco-like proteins (RLPs); however, both rubiscos and RLPs share structural folds and a common evolutionary origin^5,6^. It is thought that rubisco carboxylase activity emerged from a different pre-existing catalytic function in an RLP^2^. Throughout its evolution, rubisco has been adapted to function in all three domains of life and in multiple metabolic pathways, only some of which involve carbon fixation^7^. Nonetheless, the vast majority of studies have focused on characterizing rubiscos from clades that are involved in autotrophy (*i.e.*, form I and form II); yet, this represents only a fraction of the full diversity of rubiscos found across the entire tree of life. Thus, our understanding of rubisco is largely biased and there remains large gaps in our understanding of rubisco diversity as a whole.

Beyond sequence diversity, rubisco has also emerged as a structurally diverse enzyme family that can adopt a wide range of homo- and hetero-oligomeric states. All rubiscos share a common fold with a unique N-terminal domain followed by a TIM barrel and a short C-terminal region^1^. The fundamental functional unit of rubisco is composed of an antiparallel dimer of two large subunits (RbcL, ∼50 kDa) with a C_2_ symmetry axis and two active sites located at the intradimeric interface. Across the tree of rubiscos, a variety of higher-order assemblages of dimers have been found (L_2_)_n_, including L_2_, L_4_, L_6_, L_8_ and L_10_ as well as L_8_S_8_ hetero-oligomeric structures containing the rubisco small subunit^8,9,10^. Given this structural diversity, the functional relevance and ancestral structure of early rubiscos still remains unknown, especially due to our incomplete coverage of quaternary structure across the phylogeny.

Given their roles in photosynthetic organisms, form I and form II rubiscos have been the most extensively sampled. Recent studies have thoroughly examined the biochemical diversity across sequences in these two subclades revealing a wide range of catalytic rates in rubiscos involved in the Calvin-Benson-Bassham (CBB) cycle^11,12^. Moreover, such efforts have revealed that the fastest rubisco enzymes are among the form II clade. Notably, form I and II rubiscos represent only 10.38% of non-redundant rubisco sequence diversity **(Table 1)**. Moreover, 56.8% of sequences are found in regions of the rubisco tree that have never been assigned a name or clade, representing 62.1% of phylogenetic diversity **(Table 1)**. Ultimately, improving our understanding of rubiscos in sparsely covered regions of the phylogeny may provide unique insights into the evolution of the broader enzyme family, highlighting the need to revisit the classification and naming scheme of new rubisco subclades.

**Table 1:** A large fraction of rubiscos remain uncharacterized. Sequence and phylogenetic diversity are measured from a dereplicated phylogeny (1859 clusters) described in Figure 1 generated from all rubisco sequences in NCBI (∼87k). Sequence diversity is measured as the percentage of sequences of each subclade compared to the total number of sequences excluding RLPs (n = 435). Phylogenetic diversity is calculated as the sum of the length of all branches, and the percentage of branch length attributable to each subclade. Grey shading for form I and II rows represent rubiscos involved in the CBB cycle. Red shading represents areas of the phylogeny that are currently unexplored.

| Form (based on current nomenclature) | Sequence diversity (% of total sequences) | Phylogenetic diversity (% of branch length across phylogeny) |
| --- | --- | --- |
| form I (*sensu stricto*) | 6.88% | 5.85% |
| form II | 3.50% | 2.82% |
| form II/III | 2.75% | 2.18% |
| form IIIa | 3.73% | 3.61% |
| form IIIb | 17.5% | 15.5% |
| form IIIc | 6.29% | 5.87% |
| form III-like | 2.55% | 2.07% |
| **all others** | **56.8%** | **62.1%** |
| Total | 100% | 100% |

To fill the knowledge gaps in deep-branching, non-CBB associated rubisco sequences across the full phylogeny, we heterologously expressed, systematically synthesized, purified, and carried out biochemical studies on novel rubiscos from sparsely covered regions and undersampled clades. Our findings uncover the structural origins of the broader rubisco family and reveal previously unrecognized folds within a novel, early-branching clade of exceptionally large rubiscos. With this expanded list of biochemically characterized rubiscos, we integrate such knowledge with an updated phylogeny and propose a streamlined naming convention for rubisco cladistics in order to better accommodate current and future discoveries of new subclades.

## Results

### Generation of a comprehensive rubisco diversity library

We compiled a comprehensive set of rubisco superfamily protein sequences by querying large databases — specifically NCBI’s non-redundant (nr) and env_nr datasets. Protein sequences were clustered at increasing identity thresholds to account for the high degree of redundancy, particularly within the form I clade. At a 65% identity threshold, we recovered 1859 unique sequence clusters, representing a 14.6% increase over our last effort ^1^. Representative sequences were selected from all clusters, regardless of form, and subjected to maximum likelihood phylogenetic reconstruction **(Materials and Methods)**. Across our phylogeny, we estimate that only 8.5% of sequence clusters at 65% identity excluding RLPs have been biochemically characterized for kinetic properties. We utilized our phylogeny to guide the selection of 102 phylogenetically diverse rubisco sequences for biochemical and structural characterization, focusing largely on clades with no previously characterized representatives and/or with evolutionary significance. All 102 genes were synthesized and cloned into an *E. coli* expression plasmid that has previously been used to heterologously express and purify a wide range of rubiscos ^8,12^.

### A proposed new naming convention for the rubisco superfamily

Our tree topology of the rubisco superfamily depicts *bona fide* (*i.e.*, currently referred to as forms I through III) rubiscos forming a monophyletic clade apart from RLP **(Figure 1a)**, consistent with previous phylogenetic analyses ^1,13,14^. Similarly, we observed that the forms I and II— the only two rubisco forms currently experimentally associated with CBB Cycle— are separated by several short but well-supported internal branches, and do not resolve as sibling clades **(Figure 1a)**. Given the emerging high-level stability of the rubisco superfamily tree, we undertook a reclassification of member clades that more clearly reflects their phylogenetic positioning in recent topologies. Also problematic is that current nomenclature — particularly the designation of RLPs as “form IV”— restricts the naming of newly identified clades, resulting in increasingly complex and inconsistent labels such as IIIA, IIIB, IIIC, and III-like. To address this, we introduce a new, more easily extensible naming scheme: the traditional form I remains unchanged, while form II is reclassified as form IIα, form II/III as form IIβ, form IIIA ^15^ as form III, form IIIb^15^ as form V, form IIIc ^13^as form VI, and form III-like ^16^as form VII.

**Figure 1:**
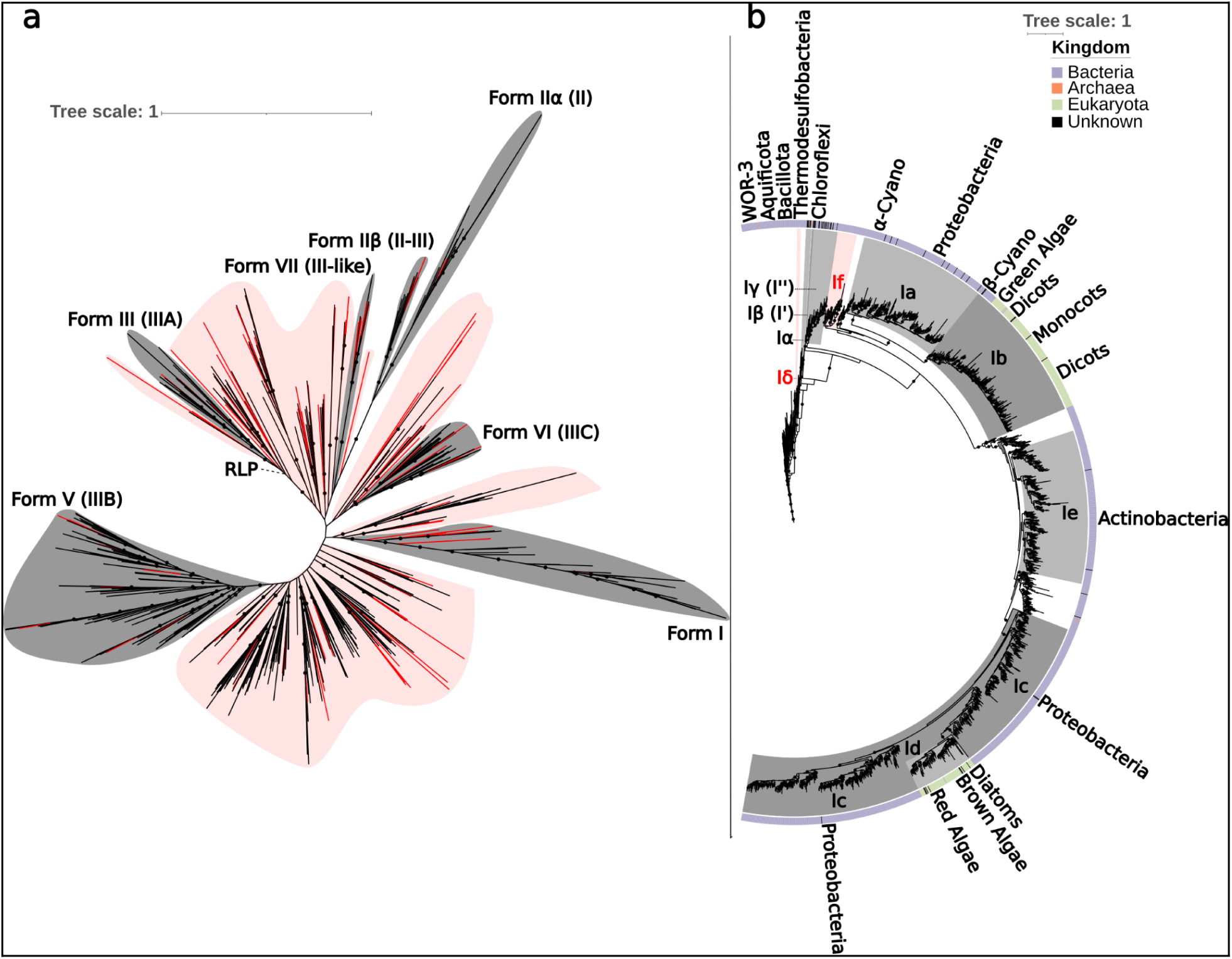
An updated phylogeny reveals many uncharacterized subclades across the rubisco protein family. **a**: phylogenetic tree of the rubisco superfamily. Highlighted in gray are known clades of rubisco labeled with their proposed clade names and current clade names in parentheses. Red branches indicate clusters that are included in our plasmid library. Black dots indicate a bootstrap value of 90% or greater. RLP is shown as a collapsed out-group. Red shading represents areas of the tree that are currently unexplored biochemically. **b**: subtree of form I clade. Highlighted in gray are known subclades of form I labeled by proposed clade names and current clade names in parentheses. A color bar around the tree indicates the kingdom that the rubisco was found in, purple: Bacteria, orange: Archaea, green: Eukaryota, gray: Unknown. Host organisms are indicated over the Form I tree but are not exhaustive. Black dots indicate a bootstrap value of >90%. Red shading newly defined clades.

Within the form I rubiscos, numerous subforms with different taxonomic, structural, and kinetic attributes are currently recognized^1^. Here, we retain the nomenclature for the previously recognized 1a, 1b, 1c, 1d, and 1e subforms **(Figure 1b)**. However, the phylogenetic relationships between these clades diverge somewhat from previous reconstructions — specifically, we now use form 1e to describe a broader, paraphyletic swath of sequences primarily associated with the phylum *Actinobacteria*, overlapping with sequence sets identified in other previous studies ^17,18^. On the other hand, we formally classify the merged, monophyletic clade form 1f. Finally, we maintain form Iα rubiscos as form 1α; however, we now refer to form I’ as Iβ, I’’ as Iγ, and formally classify a new clade Iδ. In addition to providing a clearer cataloguing of current sequence diversity, we also anticipate that the proposed scheme will facilitate easier expansion of nomenclature in the future as additional (sub)forms are discovered.

The revised phylogenetic placement of rubisco sequences provides new insight into the evolutionary trajectory of early branching form I sequences. We also observed intriguing taxonomic patterns among sequences placed within our revised form I topology. Notably, rubisco sequences at the deep base of form I — including those from members of the candidate division WOR-3 — derive from bacterial species rather than archaeal ones, distinguishing them from form V; this contrasts with the placement reported by Zeng *et al*.^19^. These basal bacteria also fall outside the Chloroflexi-associated lineages that dominate later-diverging form I groups, diversifying the evolutionary context from which canonical form Is emerge.

The earliest branching clade within form I, designated Iδ (**Figure 1b, SI Figure 1**), includes genera such as *Ammonifex (Bacillota)*, *Thermodesulfitomonas (Bacillota)*, *Thermodulovibrio (Aquificota)*, and *Thermodesulfobium (Thermodesulfobiota)*. This clade bridges the transition from deep-rooted form I sequences with L_2_ dimer oligomerization to those associated with higher-order oligomers L_8_ in form Iβ and SSU-containing L_8_S_8_ *Chloroflexi*, and the L_8_S_8_ geometry continues through form Ie (predominantly composed of sequences from the *Actinobacteria*) into the Ic and Ia groups, which are largely composed of *Proteobacteria*. This degree of taxonomic diversity suggests an extensive history of horizontal gene transfer (HGT) involving rubisco, and possibly other components of the CBB pathway, even among a set of relatively late-evolving and structurally constrained rubiscos.

In addition to their phylogenetic significance, the environmental origin of the deep-branching form I groups offer further insights into rubisco diversification. Members of the candidate division WOR-3 inhabit anaerobic sediments, hydrothermal vents, and hot springs, typically under acidic to neutral conditions (pH 2–7) and moderate to high temperatures^19,20^. These organisms are generally facultative anaerobes, engaging in hydrogenotrophic and sulfur-cycling metabolisms ^19,20^. In contrast, species within the Iδ clade — *Ammonifex, Thermodesulfitomonas, Thermodulovibrio, and Thermodesulfobium* — are strict anaerobes that thrive in thermophilic, often geothermal or subsurface environments ^21–24^. *Ammonifex* and *Thermodesulfobium* are chemolithoautotrophs, utilizing hydrogen and reducing sulfur compounds, while *Thermodesulfitomonas* and *Thermodulovibrio* display organotrophic sulfur-reducing metabolisms^25,26^. Despite broad similarities in temperature and redox conditions, WOR-3’s broader pH tolerance and facultative anaerobiosis distinguish it ecologically from its counterparts with Iδ rubiscos.

Although the current rubisco naming scheme has long served as a community standard, the rapid influx of new sequences is reshaping our understanding of rubisco phylogeny and overall topology in ways that the existing framework cannot easily accommodate. The current system lacks flexibility for introducing newly defined clades and subclades. To address this, we propose retiring the “form IV” designation in favor of referring to these sequences as RLPs, thereby freeing the nomenclature to incorporate future clades (*e.g.*, forms V, VI, VII, etc.). The goal of this updated scheme is to maximize flexibility while encouraging community engagement in recognizing clades that have strong phylogenetic support and some degree of biochemical, genetic, or physiological characterization. We anticipate lively discussion and iterative refinement of this system, much like what has occurred with the CRISPR-associated (Cas) protein family. Importantly, establishing a consistent and adaptable nomenclature will ensure that relevant clades can be clearly identified, studied, and discussed across the community, thereby accelerating scientific progress.

### Rubisco genomic context recapitulates major phylogenetic forms

In prokaryotes, rubisco genes are often found in the genome co-locating with a variety of functionally related genes like rubisco activases and other chaperones, regulators, genes involved in the synthesis of carbon concentrating mechanisms, and those involved in the CBB Cycle ^27,28^. As an alternative approach to summarizing rubisco diversity and exploring its functional associations, we undertook a genomic context analysis of diverse rubiscos drawn from our non-redundant sequence set. For each rubisco locus with adequate genomic sequence context, we retrieved all protein sequences within 10 open reading frames (ORFs) upstream and downstream, clustered neighboring proteins into families, and then performed multiple dimensionality reduction steps to determine similarities between loci based on their patterns of gene presence/absence **(Materials and Methods)**. Such an approach, previously applied to other microbial genes involved in chlorine cycling^29^, permitted the visualization of 5,774 sequence clusters sharing similar (and thus, arguably, functional) context at a high level across phylogeny/taxonomy.

Most strikingly, rubisco loci across the tree of life are organized into distinct clusters that align with their phylogenetic form, even though they share some neighboring functional genes. When sampled at a higher identity threshold (95%), we observed numerous form I associated clusters from the Ia-Ie subclades. However, genomic context could not be reliably retrieved for sequences assigned to the Iγ clade. (**SI Figure 2c**). Form I loci are mostly, though not entirely, distinct from loci with form IIα rubiscos from bacteria. Loci with form IIβ, III, V, VI, and VI formed one large, dense, cluster alongside several smaller ones associated with form IIα rubiscos (**Figure 3c**), suggesting that these rubiscos are likely functionally similar despite substantial phylogenetic divergence (**Figure 3c**). Alternatively, divergence from more canonical form I and form IIα loci could drive clustering of diverse form IIα-VII rubiscos.

**Figure 2:**
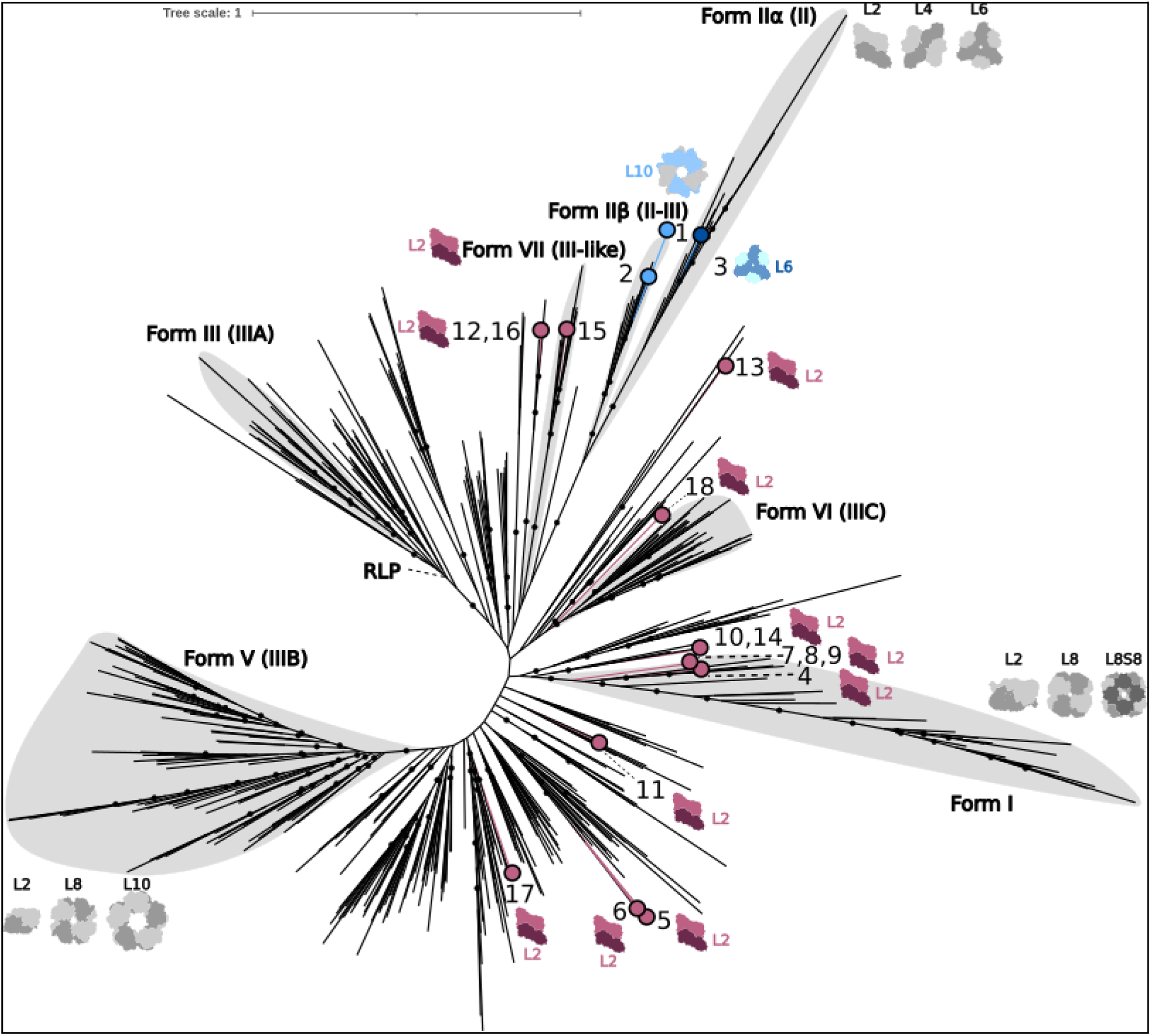
Deep branching rubiscos are dimeric. Phylogenetic tree of rubisco superfamily with oligomeric state of our 18 characterized rubiscos mapped on top. Highlighted in gray are known clades of rubisco labeled by new clade names and old clade names in parentheses. Pink branches indicate clusters that were characterized for oligomeric state. RLP is shown as a collapsed out group. Rubisco sequences are derived from the following genomes or metagenome assembled genomes. 1: *Candidatus Gottesmaniibacteriota*, 2: *Candidatus Dojkabacteria*, 3: *Rhodopseudomonas faecalis*, 4: *unclassified Bacteria*, 5: *Patescibacteria sp. (*MBU2523915.1), 6: *Patescibacteria sp. (*MBU1445715.1), 7: *Fervidicoccus fontis*, 8: *Thermodesulfobium acidiphilum*, 9: *Ammonifex degensii KC4*, 10: *Aciduliprofundum boonei*, 11: *Altiarchaeales archaeon*, 12: *Lokiarchaeota archaeon*, 13: *unclassified bacteria*, 14: *Methanobacteriota archaeon*, 15: *Kerfeldbacteria bacterium*, 16: *Prometheoarchaeum syntrophicum*, 17: *Candidatus* “Woesearchaeota archaeon CG07_land_8_20_14_0_80_44_2”, 18: *unclassified bacteria*

**Figure 3:**
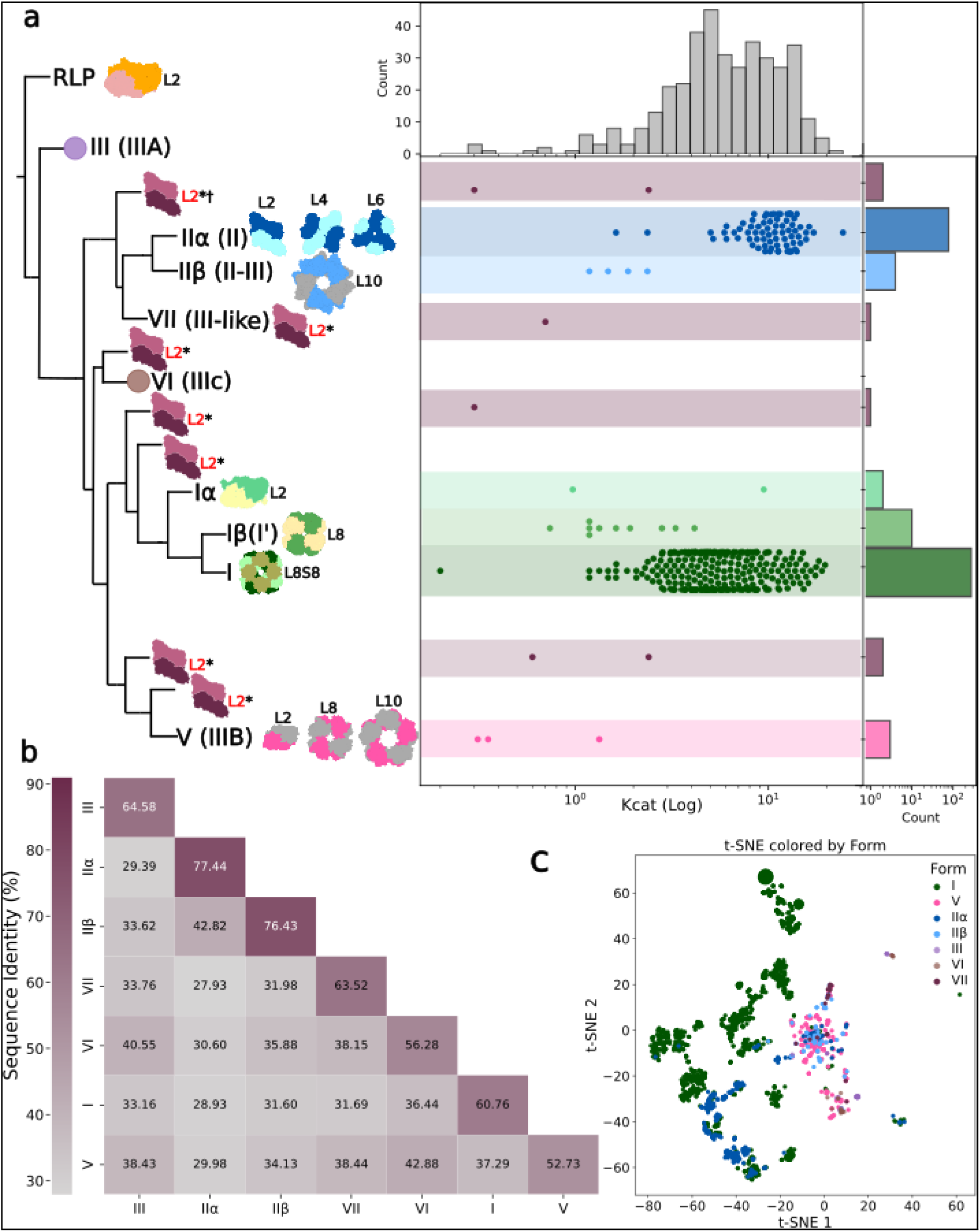
Biochemical, sequence similarity, and genomic context across the rubisco superfamily. **a:** Newly characterized oligomeric states are marked by a * and purple/pink dimer cartoon, † denotes crystal structure presented in this paper, red text denotes rubisco that are in previously unexplored regions of the tree. Known clades are labeled with proposed nomenclature and current nomenclature in parentheses. Clades or clusters with *k_catC_* (Log) measurement in line with the tree. Top histogram shows the count and distribution of *k_catC_*(Log) measurements by *k_catC_*, the right histogram shows the count and distribution of *k_catC_*(Log) measurements by form. **b:** pairwise sequence identity heatmap. Sequence identity is shown for all forms of rubisco with numeric values and color **c:** t-SNE of rubisco sequences colored by form.

Regardless, biochemically characterized rubiscos fell, as expected, primarily within form I clusters (**SI Figure 2b**); however, we also observed numerous functional/contextual clusters without any characterized representatives. Among these clusters were those associated with the form I and IIα but do not encode phosphoribulokinase (*prk*), a key gene that regenerates the rubisco substrate ribulose-1,5-bisphosphate (**SI Figure 4b**). While genes encoding the CBB Cycle are known to be spatially dispersed along the chromosome in some organisms – particularly obligate chemoautotrophs that need not tightly regulate the expression of carbon fixation ^28^– we nonetheless propose that form I or IIα sequences in clusters lacking *prk* may be novel in terms of their functionality and/or regulatory contexts, and thus should be prioritized for future study. Using comparative genomic context analysis across thousands of rubisco loci, we uncovered distinct, functionally coherent gene neighborhoods that correlate strongly with rubisco form and phylogenetic domain (**FIgure 3c**).

### The majority of deep-branching rubiscos are dimeric

From our synthesized library of 102 diverse rubisco sequences, all were heterologously expressed and purified from *E. coli*, with only 18 that yielded soluble purified enzyme for downstream biochemical characterization and investigation of their oligomeric state. Prior to this study, only four rubiscos have ever been purified from archaea. Here, we purified and characterized seven additional archaeal rubiscos, expanding representation of rubiscos in a poorly sampled domain of life^9,30–32^. Until recently, rubisco dimers were regarded as anomalous within the broader landscape of rubisco oligomerization, primarily associated with the form IIα clade ^8^. However, the discovery of form Iα and Iγ clades positioned dimeric rubisco basal to the form I lineage, suggesting a dimeric origin for the form I clade^33^. Despite these insights, the prevalence of dimeric rubisco across the full phylogenetic spectrum — particularly among newly described deep-branching clades — remained poorly understood.

To address this gap, we characterized all 18 soluble rubisco variants in our library using SEC–SAXS. This technique provided bound and unbound scattering profiles for each variant, enabling precise oligomeric state determination (**SI Figures 7, 8**). Remarkably, 15 of the 18 rubiscos analyzed — originating from previously uncharacterized regions of the phylogenetic tree — exhibited dimeric quaternary structures (**Figure 2, SI Tables 1, 2**). These diverse sequences underscore the widespread distribution of the dimeric architecture across rubisco phylogeny. Moreover, these findings reinforce the hypothesis that dimerization represents the ancestral state of rubisco.

**Table 2:**
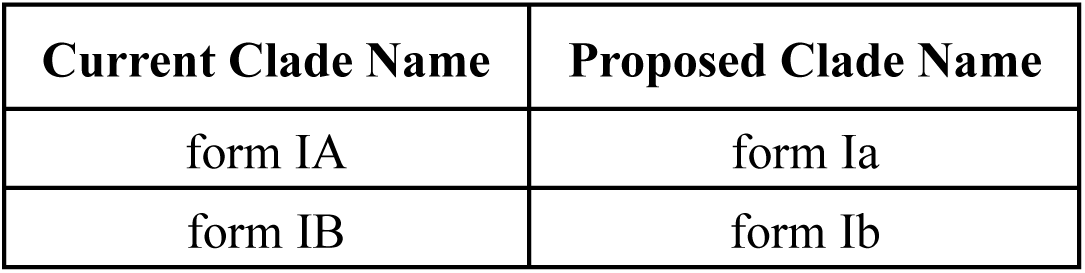
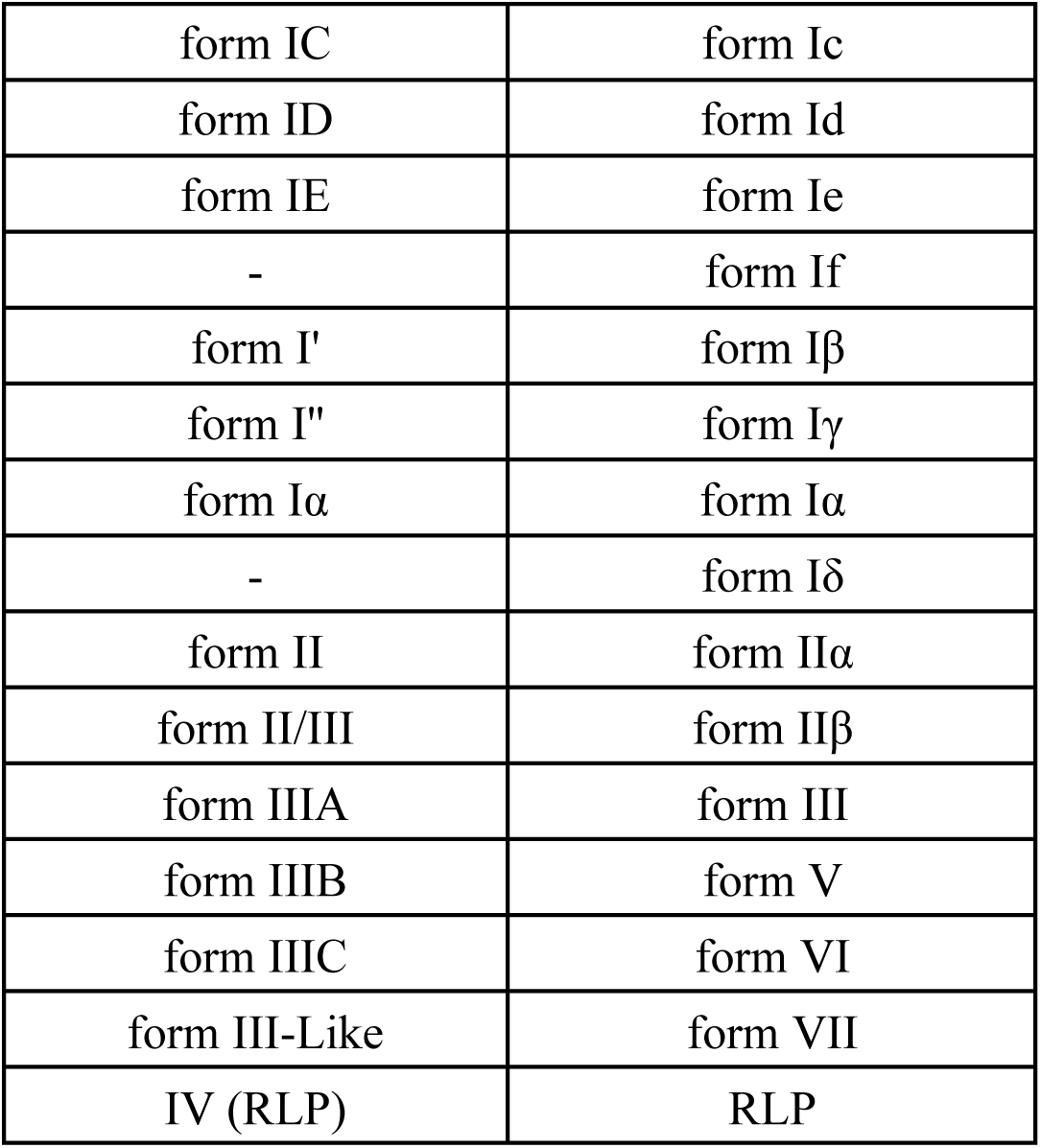
New naming convention for rubisco superfamily. Clades are denoted with Roman numerals. Subclades are denoted with the lowercase Latin alphabet if a subclade includes a small subunit, and the Greek alphabet if a small subunit is not present. form IV was excluded to avoid confusion with RLPs

In this context, higher-order assemblies (L₄, L₆, L₈, L₁₀) outside of form I appear to have arisen secondarily, likely as adaptive responses to stability demands, with oligomerization emerging as a consequence rather than a primary functional requirement. Such oligomeric complexity may have been driven by selective pressures that also shaped the evolution of the small subunit and the recruitment of protein chaperones, as strong selection for catalytic efficiency can destabilize protein structure, fostering alternative folding pathways and promoting interactions with stabilizing partners^34,35^. Structural instability — particularly under diverse environmental conditions — can lead to misfolding or aggregation, which proteins must counteract to remain viable^36^. In this evolutionary context, the prevalence of dimers among deep-branching lineages suggests that the ancestral rubisco functioned without the reinforcement of additional subunits, with higher-order oligomers evolving later as compensatory adaptations that stabilized otherwise vulnerable assemblies. In some lineages these adaptations became fixed, locking in complex architectures; in others, relaxed constraints or alternative solutions permitted reversion to the simpler, dimeric form^8,10^. This pattern underscores the dimer as the evolutionary starting point from which present-day structural diversity in rubisco arose.

Taken together, our survey of a 102-sequence library — yielding 18 soluble enzymes for SEC–SAXS analysis — shows that 15 variants from deep-branching, previously uncharacterized clades adopt dimeric quaternary structures, supporting a dimeric ancestral state and a broader prevalence of dimers than previously recognized. Within this framework, higher-order L7 assemblies are most parsimoniously interpreted as derived states that emerged in response to structural stability pressures. These findings refine models of rubisco oligomeric evolution and provide a robust, data-driven baseline for subsequent structural and mechanistic exploration.

### Biochemical characterization of phylogenetically diverse rubiscos

To complement our structural analyses, we performed biochemical characterization on a subset of phylogenetically diverse rubiscos. Thermostability (T_m_) measurements were obtained for 10 variants, while catalytic turnover rates (*k_catC_*) could only be determined for 7. Thermostability measurements were attempted for all 18 rubiscos; however, sufficient purity was achieved for only the 10 presented here. These sequences span deep-branching regions of the rubisco tree, offering a rare biochemical lens into lineages that remain largely uncharacterized.

Oligomerization has been proposed as an evolutionary strategy for enhancing protein thermal stability^37–39^. To test this hypothesis in the context of rubisco, we surveyed the literature via NCBI PubMed and identified 14 reported T_m_ values: nine from form I rubiscos, one from the form Iβ rubisco of *Candidatus Promineofilum breve*, two from form IIα rubiscos, one from form IIβ, and two from the form V rubisco. These measurements are biased toward form I enzymes and were obtained using heterogeneous methods. To broaden the scope of thermal profiling and investigate the structural constraints governing rubisco thermostability, we experimentally determined T_m_ values for 10 phylogenetically diverse rubiscos from our curated library (**SI Table 3**). Using a fluorescence-based protein thermal shift assay, we observed melting temperatures ranging from 57.5°C to 95.2°C (**SI Figure 5**).

Previous studies have shown that reducing oligomeric state—by mutating dimer–dimer interface residues in higher-order assemblies (*e.g.*, L_10_, L_6_)—can lead to decreased thermal stability, supporting a potential link between oligomerization and thermostability ^8,30,39^. However, our data from phylogenetically diverse rubiscos revealed no such correlation between oligomerization and thermostability (**SI Figure 6a**). Notably, we identified three dimeric rubiscos within the form Iδ clade that exhibited high T_m_ values ranging from 77.5°C to 95.2°C, all derived from thermophilic organisms (**SI Figure 6a**). This observation echoes findings from Bundela *et al*., who reported a dimeric rubisco with the highest known T_m_ (114.5°C), also from a thermophile^30^. In contrast, we observed a hexameric form IIα rubisco with a T_m_ of 57.6°C—comparable to the lowest recorded values (**SI Figure 6a**). Taken together, these results suggest that oligomeric state alone does not correlate with thermostability, prompting consideration of alternative factors—such as sequence composition, surface chemistry, or environmental adaptation—that may underlie thermal resilience in rubisco.

Rubisco’s catalytic efficiency has long been studied, largely in the context of its importance in photosynthesis and carbon fixation; however, kinetic parameters remain sparsely documented outside form I and IIα clades for this reason. The few enzymatically characterized non-form I and IIα rubiscos, tend to have slow *k_catC_*, yet to what extent this trend holds across non-CBB associated rubsicos has not traditionally been well studied due to poor sampling of the other clades – only recently has there been more interest in better understanding this point ^40^. To broaden this dataset, we quantified carboxylation turnover *k_cat_*_C_ for seven deep-branching rubiscos spanning underexplored phylogenetic diversity (**SI Table 4, Figure 3a**). These newly characterized enzymes include one form IIα rubisco, one from the previously unsampled form VII clade, and five additional variants from uncharted regions of the rubisco tree.

To contextualize rubisco’s catalytic constraints, we compared turnover rates across newly characterized variants, integrating our measurements with recent efforts to expand kinetic profiling beyond historically overrepresented clades (**Figure 3a**). Previous studies have highlighted potential sampling bias toward slower, predominantly eukaryotic rubiscos, suggesting that true median activity may be underestimated ^12,40,41^. A recent expansion of kinetic measurements across forms I, Iα, IIβ, and V supports an updated global median of ≈6 s⁻¹, approaching that of typical enzymes measured at ≈10 s⁻¹ ^40,42,43^. Yet rubisco remains catalytically constrained relative to central metabolic enzymes, whose *k_cat_*_C_ values often range between 20–80 s⁻¹^42,43^. Our data reinforce this trend while adding new kinetic measurements from underrepresented rubisco clades, contributing to a more complete view of rubisco’s catalytic landscape. Notably, two of the variants we characterized — *Prometheoarchaeum syntrophicum* and *Patescibacteria* sp. (*MBU2523915.1)* — exhibit relatively fast turnover rates (k_catC_ = 2.4 s⁻¹), compared to nearly all deep-branching, non-CBB related rubiscos.

### Large, deep-branching rubisco highlights structural diversity within dimers

Beyond phylogenetic diversity, the expanding catalog of rubisco sequences reveals variants with striking structural features. Two proteins — *L. archaeon* and *P. syntrophicum* — identified in a previously uncharacterized deep-branching clade basal to form VII, stood out due to their exceptional sequence lengths: 557 and 559 amino acids, respectively. These are approximately 13% longer than any rubisco characterized to date and 20% larger than the average rubisco length of 466 amino acids (**Figure 4**). Their extended sequences include expanded N- and C-termini and multiple novel insertions, representing primary structure deviations not observed in any other rubisco (**Figure 4a, 4c**).

**Figure 4:**
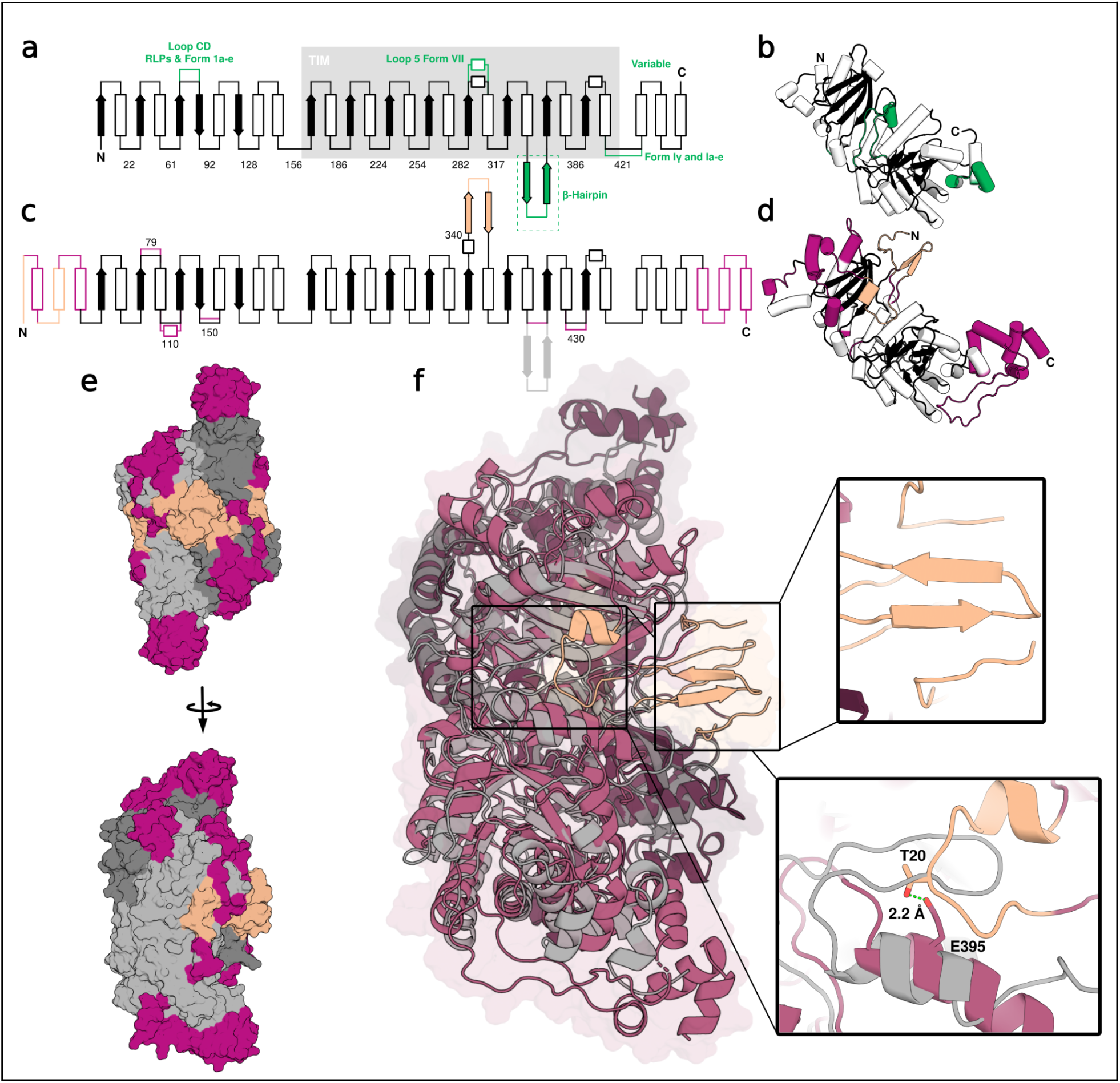
Novel, deep-branching rubisco reveals unprecedented structural diversity of the core rubisco dimer. **a:** 2d topology of rubisco consensus, **b:** tertiary structure of consensus rubisco **c:** 2d topology of *Prometheoarchaeum syntrophicum* rubisco. Insertions shown in purple, with β-sheet and N-terminus shown in orange. **d:** tertiary structure of *Prometheoarchaeum syntrophicum* rubisco, Insertions shown in purple, with β-sheet and N-terminus shown in orange. **e:** surface representation of the crystal structure of rubrum (9RUB) and apo *Prometheoarchaeum syntrophicum* rubisco (9Z01) superimposed shown in grey, and purple and pink respectively with 90° rotation. Orange as in **d**. **f:** Transparent surface and cartoon representation of apo *Prometheoarchaeum syntrophicum* rubisco structure shown in purple and pink, 9RUB shown superimposed in gray, zoom in of novel β-sheet motif shown in orange, and missing β-hairpin and functional equivalent loop light gray and orange, respectively. 2.2Å sidechain-sidechain hydrogen bond shown with green dashed line.

The first insertion, spanning residues 79–84, modestly elongates the loop connecting β-strand 2 to α-helix 5. A second insertion, located at positions 108–124, introduces a short α-helix (helix 6) that bridges α-helix 5 and β-strand 3. Most strikingly, a third insertion between residues 340–354 forms a completely novel β-sheet through interaction with the terminal region of the N-terminus—an architecture not observed in canonical rubiscos. In addition to these insertions, we identified a major deletion (**Figure 4c**): the absence of a β-hairpin motif typically conserved in *bona fide* rubiscos^2^. This hairpin, which extends prominently in forms IIβ and VII, is notably absent in RLPs and missing in these deep-branching variants.

These changes in primary and secondary structure cascade into the tertiary structure, resulting in visibly distinct monomeric folds compared to the consensus model or any previously characterized rubisco (**Figure 4b, 4d**). Together, these structural differences suggest functional divergence and raise intriguing questions about the evolutionary pressures shaping rubisco architecture in these lineages — a hypothesis further supported by the low carboxylation turnover rates observed for this variant.

To explore structure–function relationships in deep-branching rubiscos, we determined the crystal structure of *P. syntrophicum*, a representative of this unclassified clade with a k_catC_ of 2.4 s⁻¹ (**SI Table 4**). The structure confirms multiple novel secondary-structure motifs observed in the sequence analysis. Most notably, the canonical β-hairpin motif following loop 6 — typically present in bona fide carboxylating rubiscos and essential for active-site stabilization — is absent. Strikingly, *P. syntrophicum* appears to compensate for this deletion through a novel interaction: residues 16–26 form a novel extended loop that engages in a 2.2Å side-chain–side-chain hydrogen bond between T20 and E395 (**Figure 4f**). This interaction stabilizes the active-site region and may substitute mechanically for the missing β-hairpin. In addition, residues 16–22 contribute hydrophobic packing that mimics the β-hairpin’s structural support. A further stabilizing element is provided by residues 342–353, which form a β-sheet that interacts with N-terminal residues 3–6 via an extended hydrogen-bonding network (**Figure 4f**). This linkage could cascade to stabilize the 16–26 loop, suggesting a convergent strategy for maintaining carboxylation competency.

Relative to *R. rubrum* (**Figure 4e, 4f**), a well-characterized form IIα rubisco, *P. syntrophicum* additionally exhibits an expanded C-terminal region comprising a short terminal α-helix bundle followed by a large unstructured loop spanning residues 502–533. This extension, in combination with the insertions described above, may influence the rigidity of the quaternary structure and, despite being distal from the active sites, could ultimately affect carboxylation efficiency. Phylogenetically, *P. syntrophicum* occupies a deep-branching position within the rubisco clade, and its atypical structural features may reflect early divergence from canonical carboxylation architectures, offering insight into the range of structural plasticity tolerated within the rubisco lineage.

Together, these findings underscore rubisco’s structural plasticity and reveal an alternative architectural solution for preserving carboxylation function in the absence of canonical motifs. The discovery of these compensatory mechanisms shows that rubisco can evolve alternative structural solutions to maintain carboxylation function when canonical motifs are lost and reinforces the need for deeper structural and functional exploration across rubisco clades. Future work to validate the functional role of these structural features will be critical for deepening our understanding of the rubisco structure–function relationship. These motifs may represent an alternative architectural solution and offer an intriguing avenue for future studies into rubisco evolution and engineering.

## Discussion

This study expands the functional and structural characterization of rubisco by exploring under-sampled phylogenetic clades and revising their evolutionary relationships. Our findings confirm that elevated catalytic rates have evolved only in select clades, while most rubiscos exhibit slower carboxylation kinetics. A previous study (de Pins et al.) identified one fast (k_catC_ > 2 s^-1^) rubisco outside of the form I and form II clades, but upon further analysis we place that enzyme at the base of Form I^40^. This high rate is therefore consistent with an evolution towards faster rates as rubisco became involved in the CBB cycle^44,45^.

By integrating oligomeric state profiling, kinetic analysis, and structural determination, we demonstrate that the basic dimeric oligomeric state is more widespread among diverse rubisco sequences than previously recognized, consistent with a dimeric origin of the rubisco superfamily. The observed structural and thermostability diversity, particularly among archaeal sequences, underscores the adaptive breadth of rubisco. The transition to higher-order oligomers, such as tetramers, hexamers, and decamers, could reflect adaptive strategies for coping with surface mutations that introduce catalytic instability, folding stress, or environmental sensitivity. This evolutionary logic — where quaternary structure adapts to preserve functional robustness — may extend to other oligomeric systems, particularly those subject to similar structural pressures. Alternatively, oligomeric state may reflect a neutral ratcheting mechanism, in which higher-order assemblies initially evolve to enhance rubisco expression and stability, are subsequently retained due to structural or regulatory constraints^36,46^. Mutations that would be tolerated in dimers become deleterious once higher-order oligomerization is established and preventing a return to the dimeric state, although reversion to a lower-order oligomeric state remains possible if the interprotein interface is stabilized primarily by hydrogen-bonding networks rather than by hydrophobic packing^8^. As Ng et al. point out, the more critical the protein, the more susceptible the organism is to purifying selection of mutations that disrupt the chaperone relationship or, in this case, the higher order oligomerization. Our phylogenetic analysis is consistent with an extension of this hypothesis, since rubiscos involved in the CBB have frequently developed higher order oligomers, it may be the case that the initial establishment of higher order oligomerization or, indeed, chaperone dependence, is more likely when selection is strong and near-neutral improvements in protein stability come with a strong fitness advantage.

Beyond oligomeric state, deeper structural analysis of a novel rubiscos can broaden our understanding of the diversity of folds and motifs that have evolved within this enzyme family billions of years. Notably, deep-branching forms such as *P. syntrophicum* exhibit atypical folds and insertions that expand the known structural repertoire of the enzyme, highlighting a broader range of architectural plasticity and structural diversity than previously recognized. The absence of the β-hairpin in this structure is only seen in forms III and VI, as well as all RLPs. The compensation of the β-hairpin in this structure (**Figure. 4f)** by an extension of Loop 5 raises the question of whether such compensation is necessary for rubisco activity. RLP structures do not compensate in this location (i.e. PDB structures 3QFW or 3FK4). Further structural studies on form III and VI may or may not reveal additional compensations testifying to the importance of this structural feature.

As this diversity continues to expand through increased sampling, the limitations of the legacy rubisco nomenclature become increasingly apparent, underscoring the need for a system that can accommodate newly-resolved clades. The legacy rubisco naming scheme, though historically important, was devised for a narrower phylogenetic landscape and struggles to accommodate the diversity now resolved across the superfamily. As clades at the base of the form Is are discovered, they shed light about the early evolution of the CBB. Similarly, as more ancient clades are resolved within the archaeal diversity, they will inform studies of the emergence of rubisco from the RLPs^2^. Our proposed framework replaces these constraints with a system that can incorporate new clades as they emerge, ensuring that terminology reflects evolutionary reality while remaining accessible to the research community.

Despite being one of the most abundant and consequential enzymes on Earth, rubisco remains unevenly characterized across its vast phylogenetic diversity, with only a fraction of lineages represented by purified and experimentally studied enzymes. This study underscores the difficulty of studying rubiscos from deep branching lineages and archaea; to date, only eleven rubiscos have been purified from an archaeal lineage, seven of which being described in this work. More comprehensive sampling of these clades, which represent the majority of rubisco sequence diversity, may yet reveal surprises. Further sampling at the base of the form I and IIα branches is also necessary to follow the evolution of rubisco as it became the primary carboxylase on our planet.

## Methods

### Phylogenetic tree inference for the rubisco superfamily

We began by building a comprehensive set of protein sequences from the rubisco superfamily, prioritizing large sequence databases and a subset of publications amalgamating rubisco kinetic data. Specifically, we queried the National Center for Biotechnology Information (NCBI) non-redundant protein database (*nr*) as well as the *env_nr* (November 2024) database using a previously published set of form-specific Hidden Markov Models (HMMs) ^47^ using hmmsearch from the HMMER suite^48^. Alignments were filtered to those in which target sequences obtained 90% or greater of overall model coverage. Resulting target sequences were appended to a separate set selected for biochemical characterization in this study, as well as those reported in previous publications ^11,12,40,41,49^. To remove partial or composite sequences, the total set was then secondarily filtered to those falling in between 358 and 716 amino acids in length.

Size-filtered sequences were next clustered using USEARCH (*-cluster_fast)* at a variety of identity thresholds. For each cluster at a 65% sequence identity threshold, we selected a representative, prioritizing those which we had attempted biochemical characterization, or, secondarily, those with the largest sequence lengths. Sequences from the Protein Data Bank (PDB) were not selected as representatives due to the frequent appearance of N- and C-terminal expression tags. Similarly, we removed a small number (n=6) of RLP with unstable phylogenetic positions in preliminary reconstructions. We used the same process to more deeply sample the phylogenetic branch including form I rubiscos, this time employing a sequence identity threshold of 90%. Form VI rubiscos were used as an outgroup in this case.

For both the full tree and form I subtree, representative sequences were aligned using MAFFT (--auto --reorder)^50^, and alignments were subsequently trimmed using TRIMAL (-gt 0.1) ^51^ to remove phylogenetically uninformative positions. Sequence alignments were manually inspected in Geneious. Maximum-likelihood trees were inferred using IQTREE2 with a model selection step and 1000 ultra-fast bootstraps (-bnni, -m TEST -st AA –bb 1000)^52^ . Clades were defined through a combination of automated and manual approaches, adjusting clade membership determined in a previous study ^1^ in consideration of 1) sequence diversity 2) the degree of bootstrap support 3) degree of taxonomic conservation and 4) prior literature. Characterization status for each representative sequence was assigned based on all members of its corresponding cluster; if any kinetic measurement had been previously made for any cluster member; that cluster was marked as successfully characterized. Similarly, if kinetic measurements had been attempted but failed, we marked clusters accordingly.

### Analysis of genomic context for diverse rubiscos

To reconstruct the genomic loci for diverse rubiscos, we created a more deeply sampled sequence set from the clustering analysis performed above (95% sequence identity). For each of these rubiscos, we queried NCBI using a custom Python script that retrieves all other upstream and downstream protein sequences on the DNA fragment from which the rubisco was originally reported (code available at https://github.com/alexanderjaffe/rubisco-diversity). Taxonomic affiliations for each locus were also obtained through this method. Only loci with at least 10 open reading frames (ORFs) upstream and downstream of the rubisco sequence were retained for further analysis.

Next, all ‘neighbor’ protein sequences were clustered into protein families using MMSeqs (--cov-mode 0)^53^. Protein families were filtered to those with 5 or more members across all rubisco loci, and loci encoding RLPs were removed. Next, we computed the similarity of rubisco loci based on presence/absence patterns of their neighboring genes. This was accomplished by adapting a previous bioinformatic workflow ^29^. Briefly, a presence/absence matrix of protein families by locus was created and subjected to Principal Component Analysis (PCA) using Python’s sklearn package. The first two PCA components were then passed to t-distributed stochastic neighbor embedding (t-SNE, also from sklearn) for additional dimensionality reduction. Finally, we visualized loci along the first two t-SNE axes as a function of their rubisco form, taxonomic affiliation, and presence/absence of selected KEGG orthologies (as determined by kofamscan) ^54^.

### Plasmids, cloning

Representative rbcL genes were synthesized by JGI (sequences available as supplementary data) and cloned into a pET28 vector with an N-terminal His14-bdSUMO tag Plasmids pSF1389 ^55^

### Expression and purification of recombinant proteins

Rubiscos were prepared by cotransforming plasmids containing His14-bdSUMO-tagged rubisco RbcL into chemically competent BL21 DE3 Star *E. coli* with pBADES/EL plasmid. Cells were grown to mid-log phase at 37 °C (OD600 ≈ 0.7 - 0.8) and overexpression of GroEL/ES was induced by the addition of 0.2% w/v arabinose and further incubation for 2 h. Cells were resuspended in fresh TB media (without arabinose) with 100 mM ITPG and shaken for 16 h at 16 °C. Pelleted cells were resuspended in pH 8.0 lysis buffer (20 mM sodium phosphate, 300 mM NaCl, 10 mM imidazole, 5% glycerol, 2 mM MgCl2) with ∼5 mM PMSF and subject to a freeze–thaw cycle at −80 °C before lysis by use of a sonicator. The soluble fraction was collected by centrifugation (15,000 RCF, 20 min). Clarified cell lysate was used to resuspend pre-equilibrated Ni-NTA resin, Ni-NTA resin was allowed to settle before the column was allowed to flow. Columns were washed thoroughly before resuspension in the bdSENP1 reaction buffer (20 mM sodium phosphate pH 8.0, 300 mM NaCl, 10% glycerol). Purified bdSENP1 was added to resuspended columns and rocked gently overnight at 4 °C to facilitate cleavage of the His14-bdSUMO tag from the target protein. Flow-through from the bdSENP1 reaction was collected. Samples were stored in 20 mM sodium phosphate pH 8.0, 150 mM NaCl, 10 mM MgCl2, 10 mM NaHCO3 at −80 °C.

### PAGE analyses

SDS–PAGE samples were prepared according to standard procedures in Laemmli Sample Buffer (Bio-rad) with 2-mercaptoethanol and heated at 98 °C for 5 min. Samples were resolved on 4-12% NuPAGE™ Bis-Tris Mini Protein Gels (Invitrogen) in 1 × Tris/Glycine/SDS buffer (Bio-Rad) and stained in InstantBlue® Coomassie protein stain (Abcam).

### Small-angle X-ray–scattering (SAXS) data collection and analysis

Small-angle X-ray-scattering (SAXS) coupled in line with size-exclusion chromatography (SEC) experiments were performed with 100 µl samples containing 1-45 mg ml^−1^ of rubisco incubated with or without CABP prepared in 20 mM HEPES-OH (pH 8.0), 300 mM NaCl, 10 mM MgCl2, 10 mM NaHCO3. SEC–SAXS data were collected at the ALS beamline 12.3.1 at the Lawrence Berkeley National Laboratory^56^. The X-ray wavelength was set at λ = 1.127 Å and the sample-to-detector distance was 2,100 mm, resulting in scattering vectors (q) ranging from 0.01 to 0.4 Å−1. The scattering vector is defined as q = 4πsinθ/λ, where 2θ is the scattering angle. All experiments were performed at 20 °C, and the data were processed as described previously^57^.

Briefly, a SAXS flow cell was directly coupled with an online 1260 Infinity HPLC system (Agilent) using a Shodex KW804 column (Showa Denko). The column was equilibrated with running buffer (20 mM HEPES-OH pH 8.0, 300 mM NaCl, 10 mM MgCl2, 10 mM NaHCO3) with a flow rate of 0.5 ml min–1. A total 90 µl of sample was separated by SEC and 3-s X-ray exposures were collected continuously during a 30-min elution. The SAXS frames recorded before sample analysis were subtracted from all other frames. The subtracted frames were investigated by R_g_ derived by the Guinier approximation, I(q) = I(0) exp(–q^2^R_g_2/3) with the limits qR_g_ < 1.6. The elution peak was mapped by comparing the integral of ratios to background and R_g_ relative to the recorded frame using the program RAW ^58^. Uniform R_g_ values across an elution peak represent a homogenous assembly. Final merged SAXS profiles, derived by integrating multiple frames across the elution peak, were used for further analysis, including Guinier plot, which determined aggregation-free state. The molecular weight from SAXS was determined by Volume of Correlation (Vc) ^59^.

All SEC-SAXS data, including processed data, are deposited in the Simple Scattering database (https://simplescattering.com/) with the ID codes listed in the SI Table 1 and SI Table 2.

### SAXS modelling

The atomistic model of rubiscos in the open conformation was prepared based on the crystal structure of the closed conformation presented in this study, with missing N- and C-terminal residues added using the program MODELLER^60^. Different extensions and compactions of the unfolded tails were built to screen conformational variability. The experimental SAXS profiles were then compared to theoretical scattering curves generated from these atomistic models using FoXS^61,62^. Theoretical scattering profiles were fitted to the experimental SAXS data. to validate solution-state conformations of rubiscos. Models of rubisco were obtained through ALPHAFOLD^63^.

### Crystallization and structural determination of rubisco

Ni-NTA–purified rubisco were further subject SEC on a Superose 6 Increase 10/300 GL, in a final buffer containing 100 mM Hepes (pH 8), 100 mM NaCl, 25 mM MgCl2, 5 mM NaHCO3, and 1 mM DTT. The dimeric rubisco Prometheoarchaeum syntrophicum (PsRuB) was screened against the following crystallization screens: MCSG-1 (Anatrace); Crystal Screen, SaltRx, PEG/Ion, Index, and PEGRx (Hampton Research); and Berkeley Screen ^64^. Crystals of the Prometheoarchaeum syntrophicum rubisco were found in 100 mM Bis-Tris-HCl pH 6.5, 100 mM Ammonium phosphate dibasic, 20 % (w/v) PEG 3,350 and 5 % (v/v/) 2-Propanol. Crystals from the *P. syntrophicum* rubisco were then placed in a reservoir solution containing 20% (v/v) glycerol and flash-cooled in liquid nitrogen.

The X-ray dataset for the PsRuB was collected at the Berkeley Center for Structural Biology beamline 8.2.1 at the Advanced Light Source at Lawrence Berkeley National Laboratory, The diffraction data were processed using the program Xia2^65^. The crystal structure of PsRuB was solved by molecular replacement with the program PHASER ^66^ using the search model generated by ALPHAFOLD ^63^. The atomic positions obtained from the molecular replacement were used to initiate model building using phenix.autobuild within the Phenix suite ^67,68^. Structure refinement was performed using the phenix.refine program ^69^. Manual rebuilding was done using COOT ^70^. Root mean square deviation differences from ideal geometries for bond lengths, angles, and dihedrals were calculated with Phenix ^68^. The stereochemical quality of the final models of *Prometheoarchaeum syntrophicum* were assessed by the program MOLPROBITY ^71^. A summary of crystal parameters, data collection, and refinement statistics can be found in SI table 5. Structures and coordinates for PsRuB can be found in the PDB under accession ID 9Z01.

### Rubisco activity assays

The rubisco carboxylation rate was measured using an NADH spectroscopic assay coupled to 3-phosphoglycerate (3-PGA) production, as previously described^12,72,73^. rubisco was activated in the presence of all assay components minus ribulose-1,5-bisphosphate (RuBP), the substrate.

RuBP was synthesised in-house as previously described ^74^. Assay activating conditions include; pH 8, 25 °C, 4% CO_2_ and 0.5% O_2_ with orbital shaking at 1440 rpm for ∼20 minutes in a 96 well flat-bottom transparent plate (Corning, Costar). The reaction was initiated with the addition of RuBP. NADH oxidation at 340 nm was used to estimate the RuBP turnover rate, accounting for the production of 2 3-PGA molecules per molecule of RuBP. Rubisco active site concentration was estimated using the known rubisco inhibitor, CABP, at n=2, using linear regression (**SI Figure 3**). Rates of carboxylation were calculated using a custom Python script (10.5281/zenodo.7757660, n=3). Two batches of CABP were synthesized from RuBP using both cold and ^14^C-labeled KCN ^75,76^. An aliquot of activated *R. rubrum* was used for determination of rubisco active sites *via* ^14^C-CABP binding and liquid scintillation, as previously described^77^. The ^14^C-CABP quantified *R. rubrum* rubisco was then used to quantify the cold CABP for use in the NADH assay.

### Protein thermal shift (PTS) assay

The PTS assay was conducted using a Protein Thermal Shift kit (Thermo Fisher). Samples were prepared with 1 mg ml^−1^ protein in 1 × PTS phosphate buffer and 4 × PTS dye in Thermo Fisher MicroAmp Optical 8-Tube Strips. The assay was conducted on an Applied Biosciences QuantStudio 3 machine for quantitative PCR with reverse transcription. The assay consisted of initial cooling and hold at 16 °C for 1 min, followed by an 0.05 °C s^−1^ increase to 100 °C and a final hold at 100 °C for 1 min. Data were analysed in Protein Thermal Shift Software.

## Supporting information

Supplemental data set 3B

Supplemental data set 1B

Supplemental data set 1A

Supplemental data set 6A

Supplemental data set 6B

Supplemental data set 5B

Supplemental data set 5A

Supplemental data set 4B

Supplemental data set 4A

Supplemental data set 2A

Supplemental data set 2B

Supplemental data set 3A

## Acknowledgments

This work was part of the DOE Joint BioEnergy Institute (https://www.jbei.org) supported by the U. S. Department of Energy, Office of Science, Office of Biological and Environmental Research, through contract DE-AC02-05CH11231 between Lawrence Berkeley National Laboratory and the U.S. Department of Energy. The work (proposal:509657 DOI:10.46936/10.25585/60008804) conducted by the U.S. Department of Energy Joint Genome Institute (https://ror.org/04xm1d337), a DOE Office of Science User Facility, is supported by the Office of Science of the U.S. Department of Energy operated under Contract No. DE-AC02-05CH11231. SAXS data was collected at the Advanced Light Source (ALS) at the SIBYLS beamline, through the Integrated Diffraction Analysis Technologies (IDAT) program, supported by DOE Office of Biological and Environmental Research. Additional support comes from the National Institute of Health project ALS-ENABLE (P30 GM124169). A.L.J. was supported by the Stanford Science Fellows program. L.M.W. was supported by the National Science Foundation Graduate Research Fellowship Program.

## Contributions

A.J.K, P.M.S, and N.P carried out overall experimental design. A.L.J, L.V.A and N.P carried out generation of a comprehensive rubisco diversity library. A.J.K, N.P, A.L.J, L.T.K, and P.M.S carried out the proposed naming convention for the rubisco superfamily. A.L.J, L.M.W, and A.J.K conducted the genomic context analysis. All protein samples were prepared by A.J.K. *E. coli* transformations, growth and expression were carried out by J.L and C.Y. A.J.K and M.H. conducted all SEC-SAXS-MALS analyses. J.L conducted all protein thermal shift experiments. L.T.K conducted all kinetic experiments. J.H.P. performed x-ray crystallography data acquisition, image processing, and structure determination. All authors contributed to writing and manuscript preparation.

