## Supplemental figures and tables for "Diversity-driven biochemical survey reveals dimeric structural origin of rubisco"

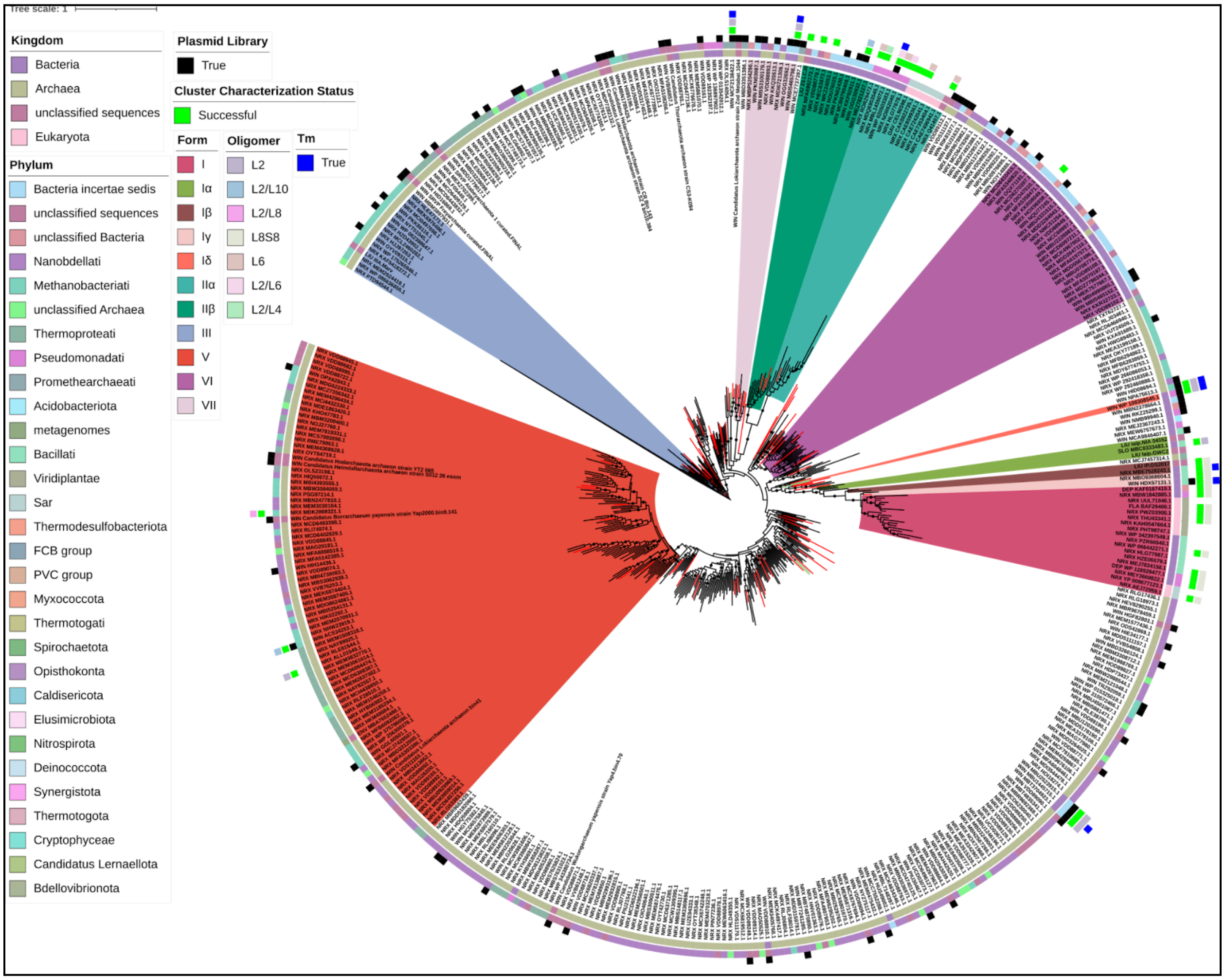


**SI Figure 1: Comprehensive phylogenetic tree highlighting rubisco clusters characterized for activity and oligomeric state**. Green highlights mark clusters previously characterized for enzymatic activity, while black bars indicate clusters analyzed in this study. Additional annotations denote oligomeric state and thermal stability (Tm).

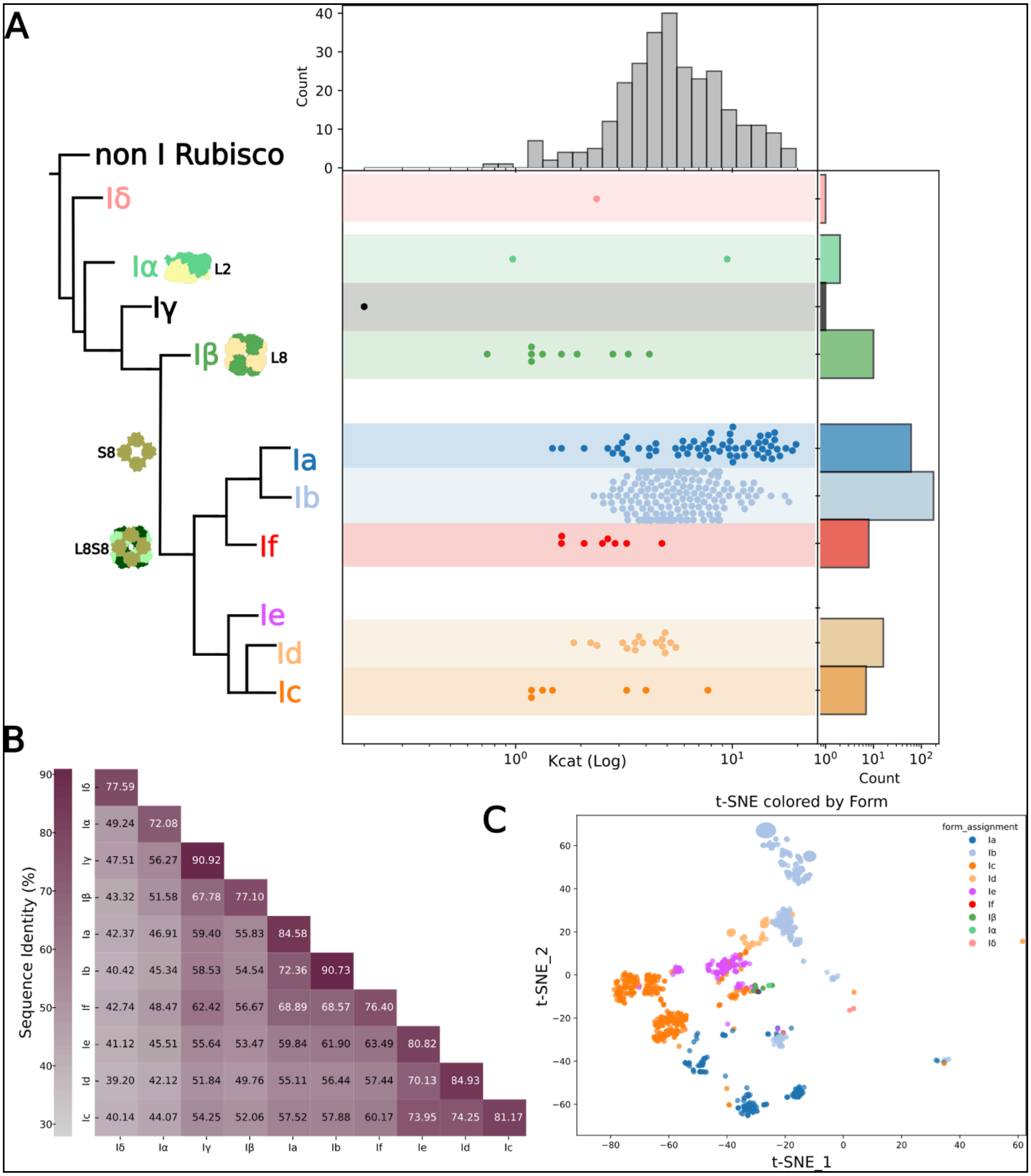


**SI Figure 2: Phylogenetic tree schematic of the rubisco form I clade; a simplified tree showing oligomeric and kinetic distribution**. **A:** Known clades are labeled with the new nomenclature. Clades or clusters with $k_{catC}$ (Log) measurement in line with the tree. Top histogram shows the count and distribution of $k_{catC}$(Log) measurements by $k_{catC}$, the right histogram shows the count and distribution of $k_{catC}$(Log) measurements by subform. **B:** pairwise sequence identity heatmap. Sequence identity is shown for all form I subforms of rubisco with numeric values and color; deep purple: >90% sequence identity, gray: ~30% sequence identity.

**C:** t-SNE of rubisco form I subform sequences colored by subform; dark blue: 1a (L8S8), light blue: 1b(L8S8), dark orange: 1c(L8S8), light orange: 1d(L8S8), purple: 1e(L8S8), red: 1f(L8S8), dark green: Iβ(L8), light green. Iα(L8), pink: Iδ(L8), black Iγ(L2).

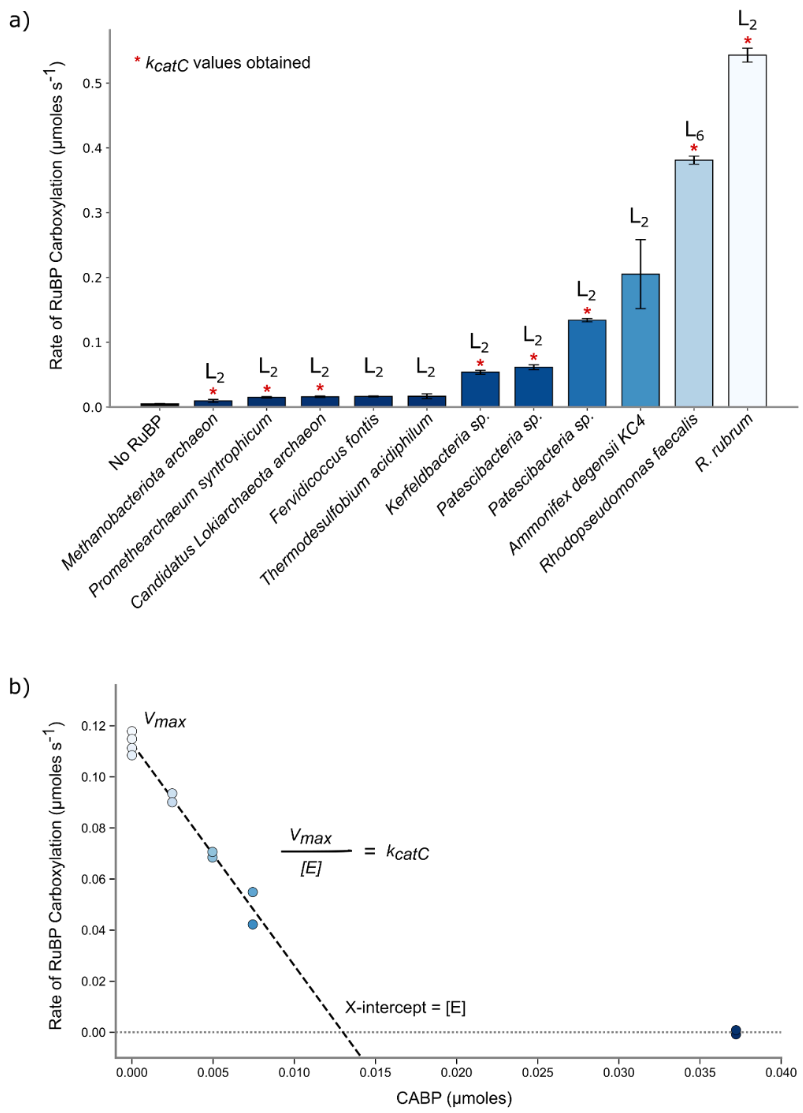


**SI Figure 3**: **Enzymatic Activity and Carboxylation Kinetics of Rubisco. a)** The activity profiles of all active rubisco sequences. Those for which $k_{catC}$ values could be determined using a CABP-titration assay are marked with *. **b)** The rate of rubisco carboxylation as a function of CABP concentration. The x-intercept gives the concentration of rubisco active sites, [E], while the y-intercept gives the reaction rate without CABP inhibition, $V_{max}$. Here, the turnover number, or $k_{cat}$ for carboxylation, of *R. rubrum* ≈ 9 $s^{-1}$.

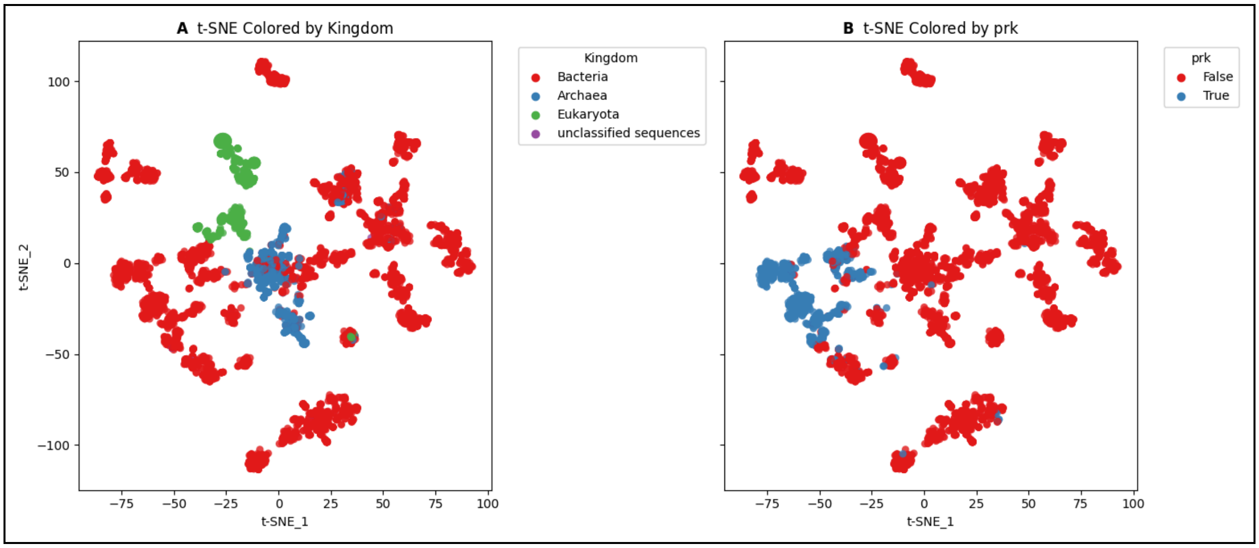


**SI Figure 4**: **TSNE color-coded by domain of life and prk .** t-SNE of all rubisco, including RLPs colored by kingdom (A) and presence of PRK (B)

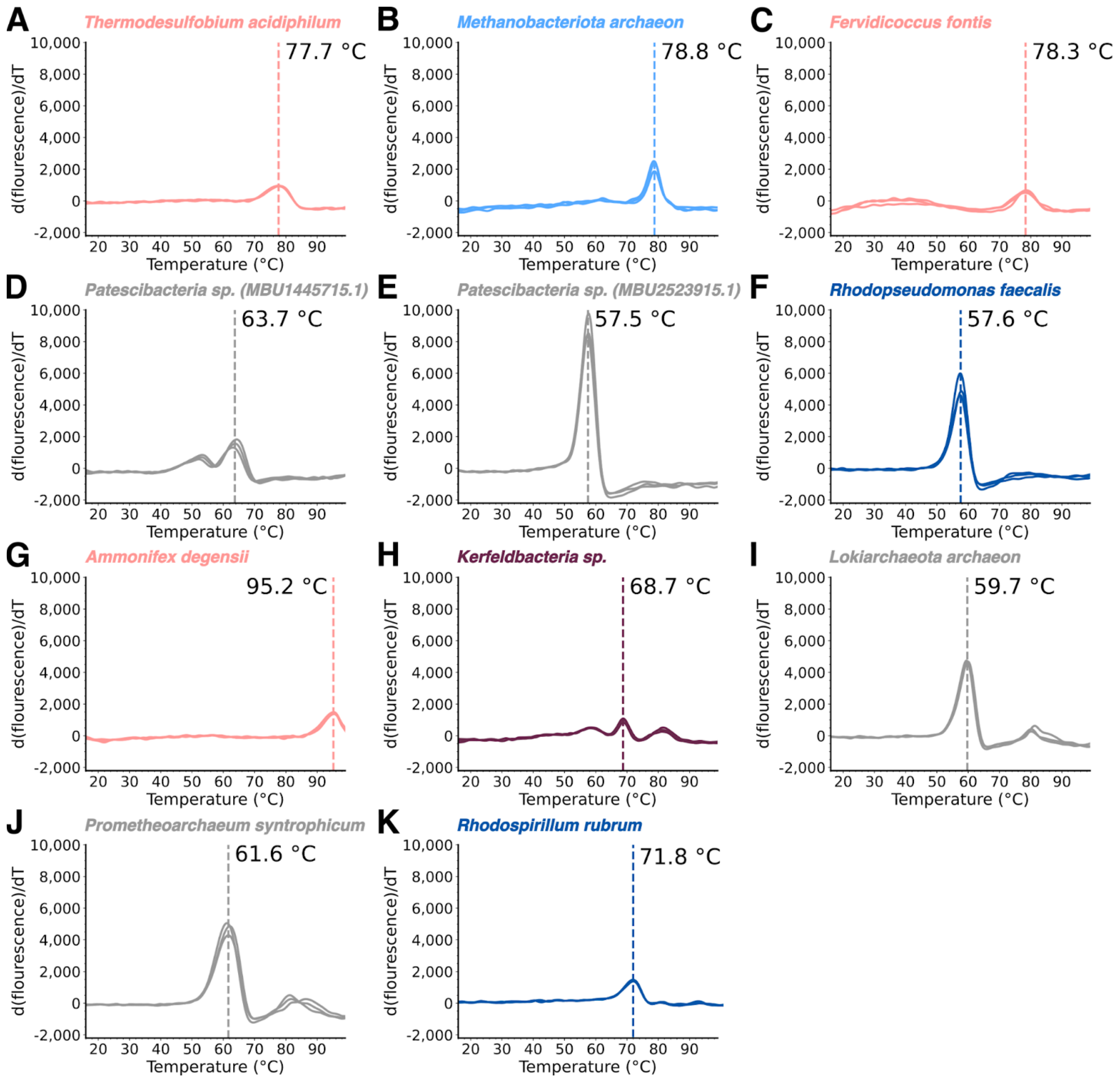


**SI Figure 5: Melting curves for rubiscos described in SI table 3. A-K)** Protein thermal shift assay data with annotated melting temperatures ($T_m$) of newly characterized rubisco dimers. Reported $T_m$ values represent the average obtained from three technical replicates.

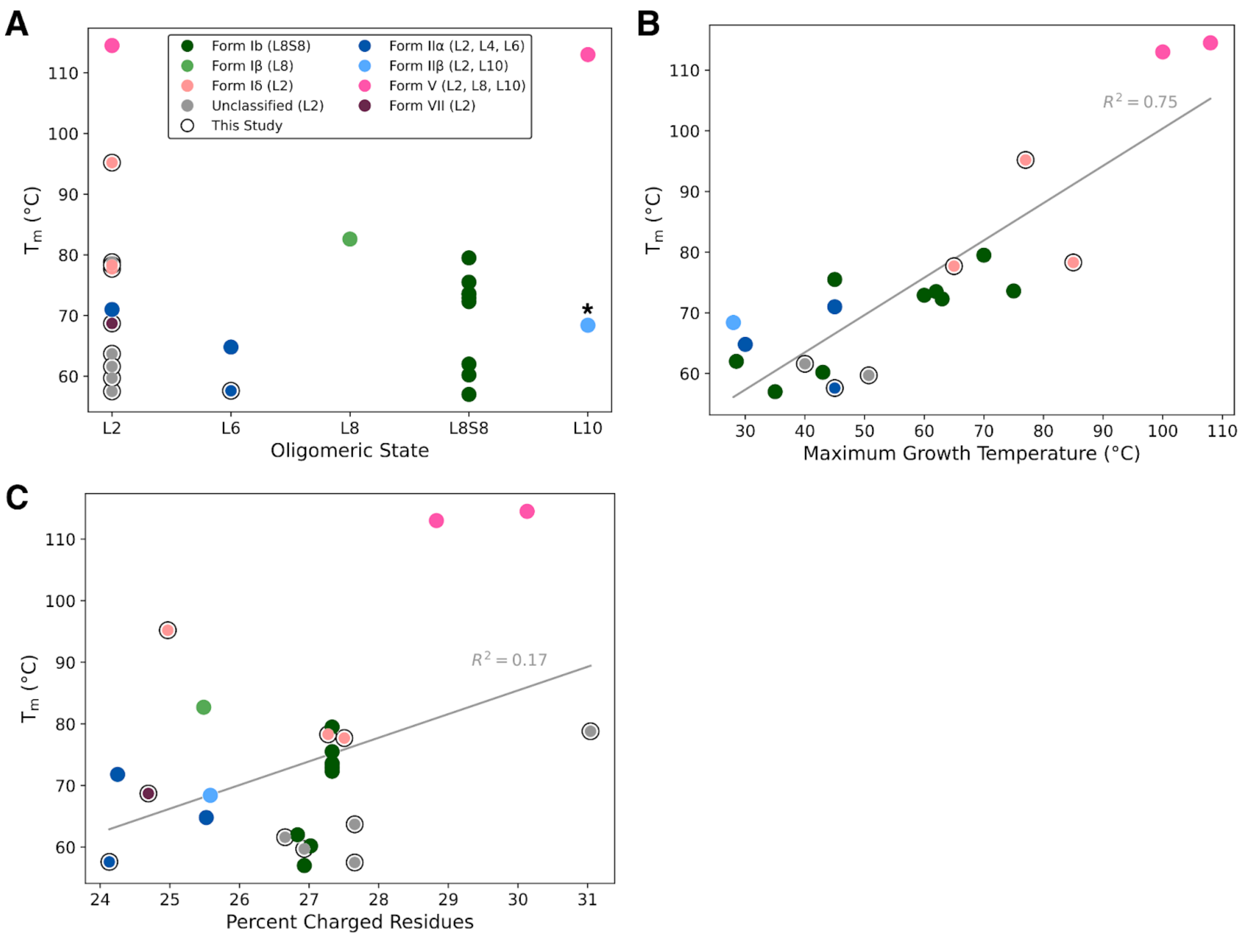


**SI Figure 6. Determinants of rubisco thermal stability: weak influence of oligomeric state and charge, strong correlation with source organism growth temperature. A)** $T_m$ plotted against rubisco's oligomeric state reveals minimal correlation. **B)** $T_m$ of rubisco plotted against the maximum growth temperature of its source organism. Positive correlation between $T_m$ and maximum growth temperature suggests that evolution of thermal stability in rubiscos was driven by the temperature of its source organism's habitat as opposed to oligomeric state. Asterisk: *Methanococcoides burtonii* can assemble into both dimer and decamer based on the presence of substrate, inhibitors or divalent metal ions. Plot reports $T_m$ of *Methanococcoides burtonii* as decamer. **C)** $T_m$ plotted against percent charged residues reveals minimal correlation.

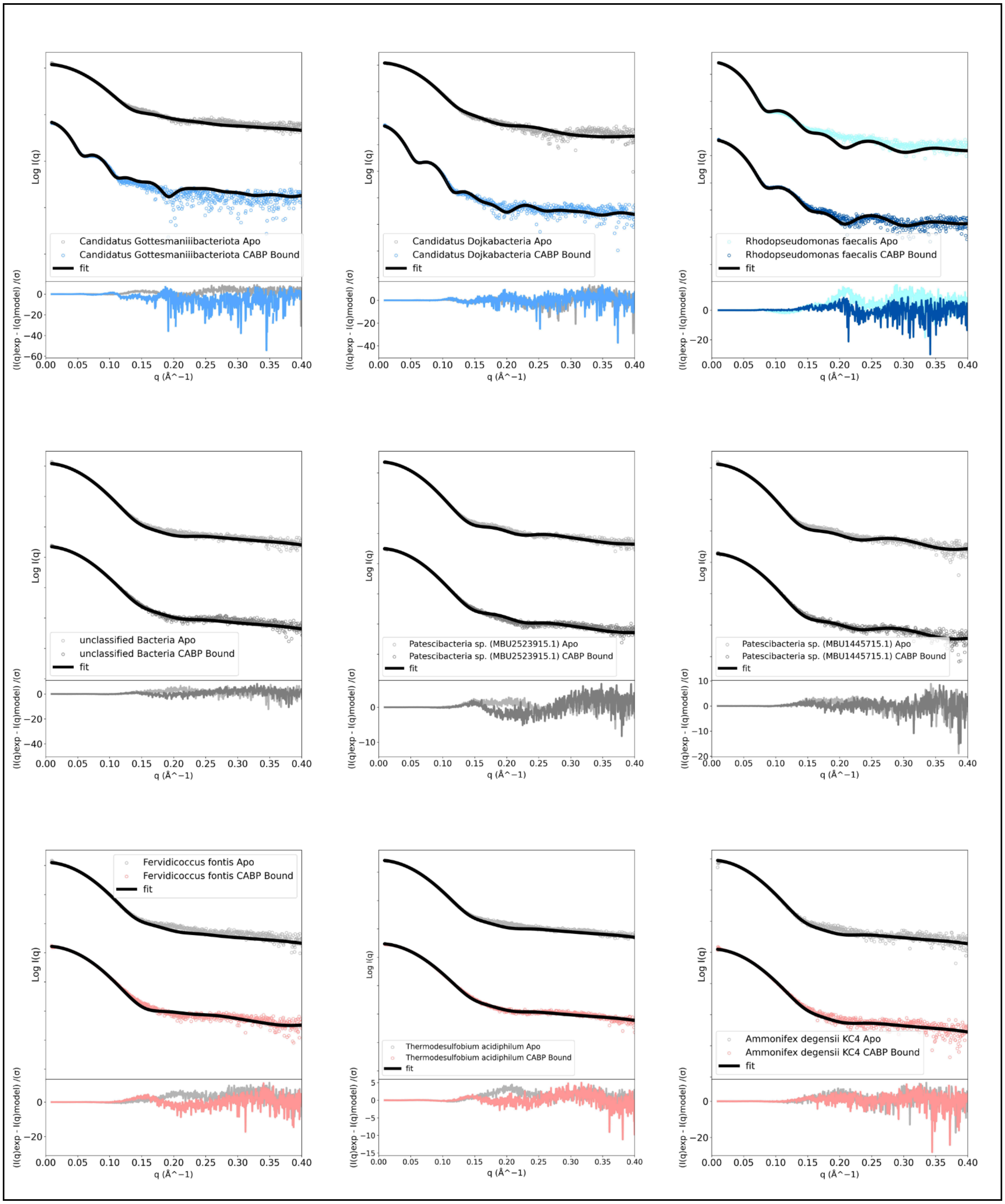


**SI Figure 7**: **SAXS scattering profiles with fitted curves for rubiscos from *Candidatus Gottesmaniiibacteriota* through *Ammonifex degensii* KC4.** Experimental scattering data are shown as colored points, with best-fit theoretical curves overlaid (black lines). Residuals for each fit are plotted below the corresponding scattering profile in matching colors. Each panel is labeled with the source organism. Together, these data provide low-resolution structural validation across a phylogenetically diverse set of rubiscos.

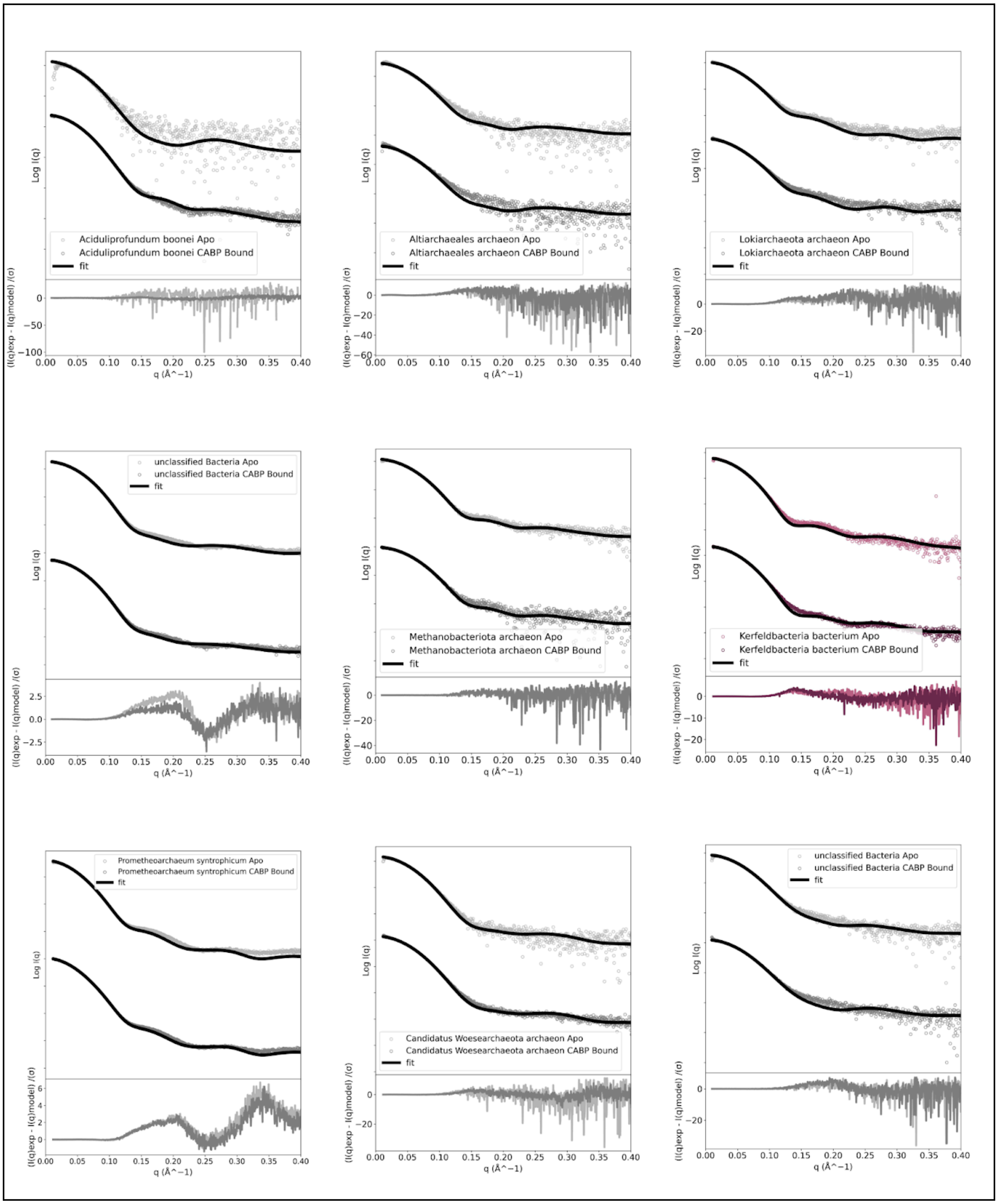


**SI Figure 8**: **SAXS scattering profiles with fitted curves for rubiscos from** ***Aciduliprofundum boonei*** **through** ***unclassified Bacteria*****.** Experimental scattering data are shown as colored points, with best-fit theoretical curves overlaid (black lines). Residuals for each fit are plotted below the corresponding scattering profile in matching colors. Each panel is labeled with the source organism. Together, these data provide low-resolution structural validation across a phylogenetically diverse set of rubiscos.

**SI Table 1: SEC-SAXS oligomeric state determination CABP bound**

| Simple Scattering Code | Organism | MW(kDa) Sequence | MW(kDa) | Rg(Å) | $\chi^2$ | Oligomeric state |
|---|---|---|---|---|---|---|
| *XSF2W1AA* | *Candidatus Gottesmaniibacteriota* | *54* | 548 | 58 | 4.4 | L10 |
| *XSZNZ3J0* | *Candidatus Dojkabacteria* | *52* | - | 56 | 2.0 | L10 |
| *XSQSTHGE* | *Rhodopseudomonas faecalis* | *50* | 261 | 43 | 1.3 | L6 |
| *XSBQGHPR* | *unclassified bacteria* | *45* | 71 | 29 | 2.6 | L2 |
| *XSGT6MLN* | *Patescibacteria sp. (MBU2523915.1)* | *50* | 79 | 29 | 2.4 | L2 |
| *XSKDNJSI* | *Patescibacteria sp. (MBU1445715.1)* | *48* | 75 | 29 | 1.1 | L2 |
| *XSXPYDB3* | *Fervidicoccus fontis* | *47* | 73 | 28 | 2.1 | L2 |
| *XS93BFEP* | *Thermodesulfobium acidiphilum* | *47* | 73 | 28 | 4.1 | L2 |
| *XSZQWWUB* | *Ammonifex degensii KC4* | *45* | - | 28 | 1.3 | L2 |
| *XSVZ5KE8* | *Aciduliprofundum boonei* | *47* | 58 | 26 | 1.0 | L2 |
| *XS895INW* | *Altiarchaeales archaeon* | *45* | 77 | 37 | 4.0 | L2 |
| *XSPQVMS8* | *Lokiarchaeota archaeon* | *62* | 97 | 35 | 6.9 | L2 |
| *XS2QFXBK* | *unclassified bacteria* | *43* | 77 | 29 | 7.8 | L2 |
| *XSFWYF0V* | *Methanobacteriota* | *47* | 74 | 28 | 1.1 | L2 |

| | | | | | | |
|---|---|---|---|---|---|---|
| | *archaeon* | | | | | |
| *XSULRK2B* | *Kerfeldbacteria bacterium* | *53* | 82 | 29 | 8.5 | L2 |
| *XSGMMWKQ* | *Prometheoarchaeum syntrophicum* | *62* | 99 | 32 | 19.8 | L2 |
| *XSMGG3DP* | *Candidatus* *"*Woesearchaeota archaeon CG07_land_8_20_14_0_80_44_2" | *47* | 75 | 29 | 4.3 | L2 |
| *XSCJOKGI* | *unclassified bacteria* | *46* | 73 | 30 | 1.4 | L2 |

**SI Table 2: SEC-SAXS oligomeric state determination unbound**

| Simple Scattering Code | Organism | MW(kDa) Sequence | MW(kDa) SAXS | Rg(Å) | $\chi^2$ | Oligomeric state |
|---|---|---|---|---|---|---|
| *XSF2W1AA* | *Candidatus Gottesmaniibacteriota* | *54* | 87 | 31 | 3.9 | L2 |
| *XSZTP5RF* | *Candidatus Dojkabacteria* | *52* | 87 | 30 | 2.2 | L2 |
| *XSPVEHHK* | *Rhodopseudomonas faecalis* | *50* | 262 | 43 | 6.4 | L6 |
| *XSKWLUDQ* | *unclassified Bacteria* | *45* | 74 | 29 | 2.5 | L2 |
| *XSPPBS9X* | *Patescibacteria sp. (MBU2523915.1)* | *50* | 84 | 30 | 2.5 | L2 |
| *XSQC2P66* | *Patescibacteria sp. (*MBU1445715.1*)* | *48* | - | 29 | 1.7 | L2 |
| *XSQ5HCCW* | *Fervidicoccus fontis* | *47* | 76 | 29 | 4.2 | L2 |
| *XSDPZBFI* | *Thermodesulfobium acidiphilum* | *47* | 76 | 29 | 6.4 | L2 |
| *XSE1JAJX* | *Ammonifex degensii KC4* | *45* | 76 | 29 | 1.9 | L2 |

| | | | | | | |
|---|---|---|---|---|---|---|
| *XSN0OSDA* | *Aciduliprofundum boonei* | *47* | 75 | 31 | 0.9 | L2 |
| *XSQIRSSH* | *Altiarchaeales archaeon* | *45* | 85 | 36 | 4.8 | L2 |
| *XSJGHHBY* | *Lokiarchaeota archaeon* | *62* | 100 | 34 | 4.6 | L2 |
| *XSEGORYK* | *unclassified bacteria* | *43* | 77 | 29 | 20 | L2 |
| *XSOXNSZO* | *Methanobacteriota archaeon* | *47* | 79 | 28 | 1.6 | L2 |
| *XSPR2OHY* | *Kerfeldbacteria bacterium* | *53* | 88 | 30 | 5.2 | L2 |
| *XSANCTQG* | *Prometheoarchaeum syntrophicum* | *62* | 101 | 32 | 37 | L2 |
| *XSDNKW9Y* | *Candidatus "*Woesearchaeota archaeon CG07_land_8_20_14_0_80_44_2" | *47* | 78 | 30 | 1.4 | L2 |
| *XSY0ANCM* | *unclassified bacteria* | *46* | 75 | 29 | 1.8 | L2 |

**SI Table 3: thermal stability of phylogenetically diverse rubiscos**

| Organism | form | $T_m$ °C | $T_m$SD |
|---|---|---|---|
| *Lokiarchaeota archaeon* | unclassified | 59.7 | 0.3 |
| *Prometheoarchaeum syntrophicum* | unclassified | 61.6 | 0.4 |
| *Patescibacteria sp. (MBU1445715.1)* | unclassified | 63.7 | 0.6 |
| *Patescibacteria sp. (MBU2523915.1)* | unclassified | 57.5 | 0.2 |
| *Methanobacteriota archaeon* | unclassified | 78.8 | 0.1 |
| *Kerfeldbacteria sp.* | VII | 68.7 | 0 |
| *Rhodopseudomonas faecalis* | IIα | 57.6 | 0.2 |
| *Rhodospirillum rubrum* | IIα | 71.8 | 0.1 |
| *Thermosulfobium acidiphilum* | Iδ | 77.7 | 0.4 |
| *Fervidicoccus fontis* | Iδ | 78.3 | 0.3 |
| *Ammonifex degensii* | Iδ | 95.2 | 0.2 |

**SI Table 4: Carboxylation activity of phylogenetically diverse rubiscos**

| Organism | form | $k_{catC}$ | $k_{catC}$SD |
|---|---|---|---|
| *Lokiarchaeota archaeon* | unclassified | 0.3 | 0.040 |
| *Prometheoarchaeum syntrophicum* | unclassified | 2.4 | 0.005 |
| *Patescibacteria sp. (MBU1445715.1)* | unclassified | 0.6 | 0.049 |
| *Patescibacteria sp. (MBU2523915.1)* | unclassified | 2.4 | 0.332 |
| *Methanobacteriota archaeon* | unclassified | 0.3 | 0.073 |
| *Kerfeldbacteria sp.* | VII | 0.7 | 0.066 |
| *Rhodopseudomonas faecalis* | IIα | 9.6 | 0.824 |
| *Rhodospirillum rubrum* | IIα | 9.8 | 0.458 |

**SI Table 5: Crystallographic statistics for X-rays data collection and structure refinement.**

| | **PsRuB** |
|---|---|
| ***Data collection*** | |
| Space group | P 1 |
| Unit-Cell parameters (Å) | 69.55 82.7 113.3 108.17 97.61 103.23 |
| Resolution range (Å) | 46.38 - 1.9 |
| Total reflections | 682393 (28461) |
| Unique reflections | 161119 (13462) |
| $R_{merge}$ (%) | 0.121 (1.15) |
| $I/\sigma I$ | 6.3 (0.9) |
| Wilson B-factor | 22.7 |
| Completeness (%) | 99.5 (98.1) |
| Redundancy | 3.8 (3.2) |
| $CC_{1/2}$ | 0.994 (0.311) |
| ***Refinement*** | |
| Resolution range (Å) | 46.38 - 1.9 (1.93 - 1.9) |
| Reflections used in refinement | 161102 (13462) |
| Reflections used for $R_{free}$ | 1809 (151) |
| $R_{work}$ | 0.1755 (0.3361) |
| $R_{free}$ | 0.2051 (0.3599) |
| No. atoms | |

| | |
|---|---|
| Proteins | 17437 |
| Ligands/ion | 40 |
| Water | 1400 |
| RMS from ideal geometry | |
| Bond lengths (Å) | 0.007 |
| Bond angles (º) | 0.71 |
| Average B-factor | 42.47 |
| Macromolecules | 42.87 |
| Ligands | 37.48 |
| Solvent | 37.60 |
| Ramachandran favored (%) | 95.99 |
| Ramachandran allowed (%) | 3.74 |
| Ramachandran outliers (%) | 0.27 |
| Clashscore | 3.36 |

[#] Values in parentheses are for the highest-resolution shell.
